# Dynamic engagement of the motor cortex in controlling movement

**DOI:** 10.64898/2026.01.13.699314

**Authors:** Tudor Dragoi, Zachary F. Loschinskey, Munib A. Hasnain, Brian DePasquale, Michael N. Economo

**Affiliations:** Graduate Program for Neuroscience, Boston University, Boston, MA; Center for Systems Neuroscience, Boston University, Boston, MA; Neurophotonics Center, Boston University, Boston, MA; Department of Biomedical Engineering, Boston University, Boston, MA; Rafik B. Hariri Institute for Computing and Computational Science and Engineering, Boston University, Boston, MA

## Abstract

Neural circuits do not contribute equally or continuously to behavior. In mice, the motor cortex can be essential or dispensable for movement in different contexts, but how it is dynamically recruited as behavioral demands evolve remains unclear. Here, we demonstrate that motor cortical involvement in movement exhibits rapid, discrete state transitions even during movements that otherwise appear continuous. While robust and reproducible across animals, the timing and presence of these state transitions are highly sensitive to task structure. We find that motor cortical engagement is sustained under conditions of sensorimotor uncertainty and dissipates rapidly when sensorimotor contingencies are resolved and actions and outcomes thus become predictable. These findings reveal a previously unrecognized layer of fast-timescale flexibility in the neural control of movement and offer a conceptual framework for understanding how cortical circuits dynamically govern behavior as demands evolve.

## INTRODUCTION

The notion of brain region specialization, which is nearly axiomatic in systems neuroscience^1–3^, implies that only a subset of all neural circuits are needed for behavior at any moment in time. It also implies that this subset evolves according to changing behavioral demands. How are different neural circuits dynamically engaged in and disengaged from behavioral control? In some circuits, such as early sensory systems, how this happens may be intuitive; primary sensory circuits are activated and may contribute to behavior whenever relevant sensory neurons are activated ^4,5^. However, in many systems that support cognition and action, theories for how different neural resources become engaged and disengaged on the timescales of behavior remain nascent^6–8^.

In the mammalian motor system, key motor centers contribute to movement in a contextually selective manner^8–11^. For example, the dorsomedial striatum supports actions that must be flexibly shaped to maximize outcome value, while the dorsolateral striatum is associated with habitual movements^11,12^. Similarly, in rodents, novel movements require the motor cortex, but the same movements may not be impaired by lesion or acute inactivation of the motor cortex once they have been learned and highly practiced^9,13–15^. During naturalistic behavior, different motor circuits are therefore presumed to engage and disengage from the control of movements dynamically, but little is known about when and how these transitions occur, and what behavioral events serve as triggers.

Here, we sought to examine when and why the motor cortex becomes engaged in – and disengaged from – the control of movement on behavioral timescales. We examined motor cortical engagement and disengagement in mice performing a suite of head-fixed licking tasks. We find not only that the motor cortex can engage and disengage from the control of movements at action transitions, but that cortical control of movements toggles on and off abruptly on faster timescales – often within individual continuous bouts of movement. Further, we find that whether and when state transitions occur can vary dramatically following subtle alterations in task structure in a manner that is highly reproducible across animals and across trials. Through a synthesis of neural population recording, behavioral and optogenetic manipulations, and statistical modeling, we demonstrate that motor cortical engagement is a function of animals’ uncertainty about future events and is consistent with a sensorimotor form of attention instantiated within the motor system.

## RESULTS

### Motor cortical activation is transient during orofacial movements

We first examined the engagement of the motor cortex during a simple task in which head-fixed animals were trained to lick a reward port for water rewards (**Fig. 1a**). In this ‘Simple Reward Task,’ head-fixed mice were trained to withhold licking during a 2-sec ‘wait’ period indicated by the illumination of an LED. Following an auditory ‘Go Cue,’ mice licked towards a single reward port that shifted randomly between one of three lateral positions every 10 trials. Mice received and consumed a water reward immediately upon their first contact (C1) with the reward port. The first lick in each bout was both longer in duration and more variable across trials than subsequent licks (two-tailed t-test, p<0.005, **Fig. 1b**; **EDFig. 1**), consistent with previous observations^16^. Mice performed hundreds of trials per session with high performance (**EDFig. 2**).

**Figure 1 -.**
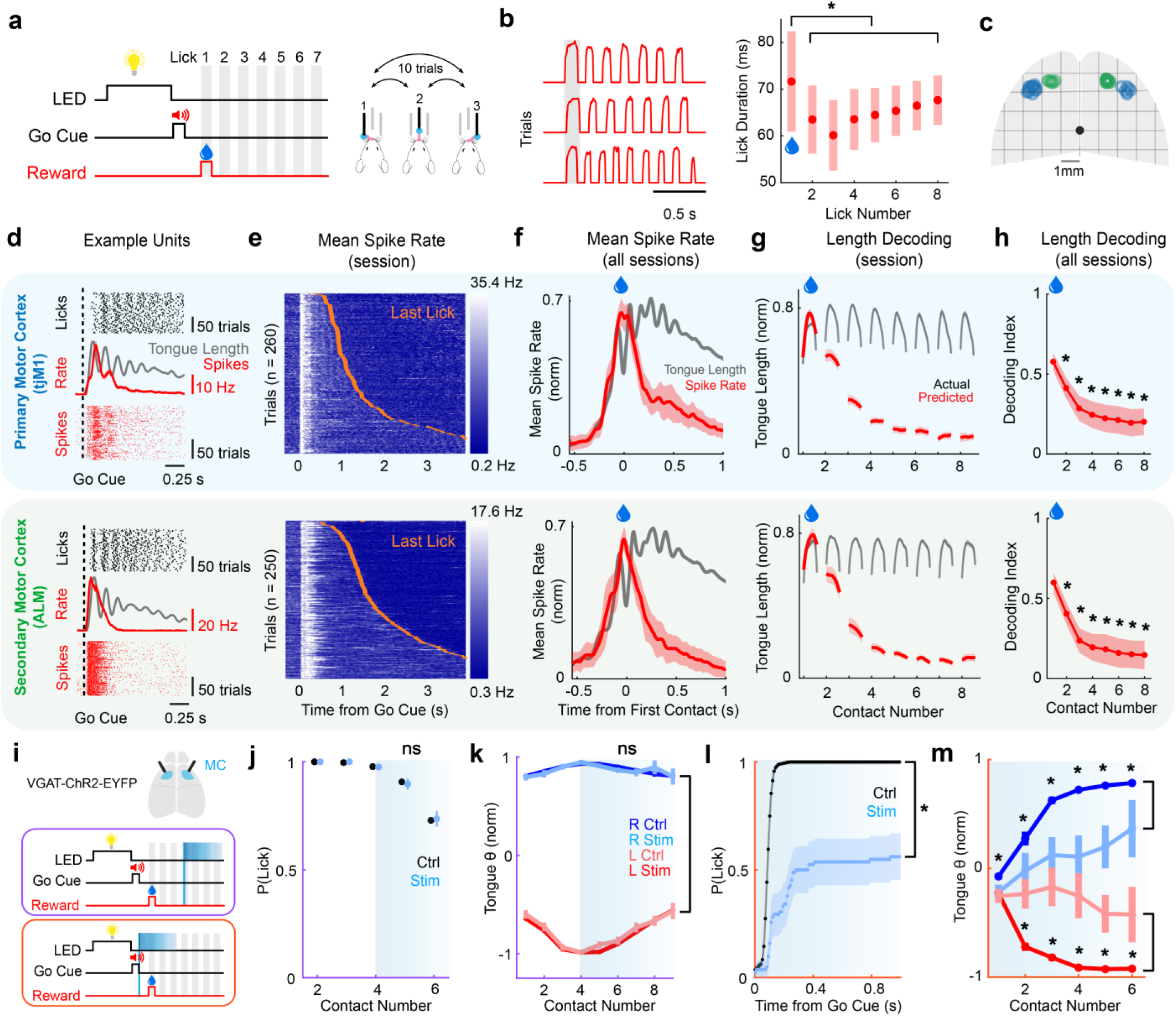
Motor cortical disengagement during the Simple Reward Task. **a.** Schematic of the Simple Reward Task. **b.** Tongue length on exemplar trials (*left*) and the mean duration of the first eight tongue protrusions across sessions (*right; n = 22 sessions, 4 animals*). The first port-contacting tongue protrusion was shorter in duration, on average, than subsequent protrusions (two-tailed t-test, p<0.005). **c.** Location of all probe insertions targeted to the ALM (*green*) and tjM1(*blue*). **d.** Rasters of port contact times (*black, top*), trial-averaged spike rate (*red, middle;* average tongue length also shown in *black*), and spike rasters (*red, bottom*) and for example units in the tjM1 (*top row*) and ALM (*bottom row*). **e.** Mean spike rate across all units in the tjM1 (*top*) and ALM (*bottom*) in an example session. Trials sorted by the timing of the last lick in a bout (*orange ticks*). **f.** Mean session-normalized spike rates in the tjM1 (*top; n = 18 sessions, 4 animals*) and ALM (*bottom; n = 12 sessions, 4 animals*) across all sessions. Average tongue length indicated in black. **g.** Tongue length predictions from the tjM1 (*top*) and ALM (*bottom*) in an example session. **h.** Decoding index (normalized model root mean squared error (RMSE), see ***Methods***) across all sessions (asterisks: p<0.005 comparing subsequent protrusions to the first, one-tailed t-test). **i.** Schematic of timing of tjM1 and ALM photoinhibition at the go cue (*orange*) or fourth port contact (*purple*). **j.** Probability of tongue protrusion on control (*black*) and fourth port contact photoinhibition trials (*light blue*; ns: p>0.005; two-tailed t-test; *n = 19 sessions, 4 animals*). **k.** Mean tongue angle on left (*red*) and right (*blue*) trials. Lighter shades represent photoinhibition trials (ns: p>0.005 comparing photoinhibition trials to control trials; two-tailed t-test). **l,m.** Same as **j,k** for photoinhibition triggered by the go cue (asterisks: p<0.005; two-tailed test comparing control and photoinhibition trials; *n = 5 sessions, 4 animals*). Shaded regions and error bars indicate 95% confidence intervals in all panels.

We recorded activity extracellularly in the tongue/jaw primary motor cortex (tjM1; n=595 single units, 18 sessions, 4 animals) and the anterolateral motor cortex (ALM; n=328 single units, 12 sessions, 4 animals), a secondary orofacial motor region, with high-density Neuropixels probes during the Simple Reward Task (**Fig. 1c**; **EDFig. 3**). Dynamics in both regions were highly modulated by movement onset, but motor cortical activation was largely transient. A large proportion of single units across both the tjM1 and ALM (48.5%) exhibited increased activity at the onset of movement, followed by a rapid decline in activity after the first tongue protrusion (**Fig. 1d**). Across all units in the tjM1 and ALM, total cortical activity consistently decayed to near baseline well before the cessation of movement (**Fig. 1e,f**).

We next evaluated the time evolution of the relationship between motor cortical activity and movement. We trained a linear decoder to predict kinematic features of movement from motor cortical activity during the initial phase of movement, when motor cortical activation was strong. The kinematics of movement, such as the cycle-by-cycle extent of tongue protrusions or the position of the jaw, could only be accurately predicted from neural activity during the first tongue protrusion (**Fig. 1g,h**; **EDFig. 4**). Decoding accuracy then decayed on a timescale similar to the overall decay in motor cortical activity. Interestingly, a small amount of residual motor cortical activation was observed to persist until movement termination. A different decoder trained on the late phase of movement, when motor cortex activation was weak, accurately predicted features of movement, such as tongue length or jaw position, throughout the duration of movement (**>EDFig. 4,5**).

To determine which components of motor cortical activity are necessary for executing movements, we optogenetically photoinhibited the tjM1 and ALM of VGAT-ChR2-EYFP mice during the early phase of movement, when motor cortical activation was robust, and during the late phase of movement, when activation was weak (**Fig. 1i**). We first applied photoinhibition after the initial strong activation of the motor cortex subsided (triggered by the 4^th^ lick contact; 10% of trials; n = 4 animals, 19 sessions; **Supplementary Video 1**). This manipulation had no detectable effect on movement kinematics. Neither the rate nor the angle of tongue protrusions (**Fig. 1j,k**) were altered, suggesting that motor cortical activity is not necessary for movement execution during the late phase. This insensitivity was observed despite the presence of residual motor cortical activation that was closely related to features of movement during the late phase. We next applied photoinhibition at movement onset (triggered by the go cue; 10% of trials; n = 4 animals, 5 sessions; **Supplementary Video 2**). In stark contrast, silencing the initial activation of the motor cortex strongly impaired the execution of movement, reducing animals’ ability to initiate movement (**Fig. 1l**). On trials where the tongue did protrude, the angle of protrusions was strongly affected such that they could not be effectively directed to lateral reward port positions (**Fig. 1m**).

Together, these results indicate a transient period of motor cortical ‘engagement’ in which it is robustly activated and necessary for the execution of tongue movements. Hundreds of milliseconds later, during the same bout of movement, activity largely dissipates, and the execution of tongue movements is insensitive to motor cortical photoinhibition. The residual activity during the late phase of movement may therefore reflect, for example, persistent sensory feedback or prediction signals rather than dynamics necessary for executing movements.

### Delaying reward prolongs cortical engagement

The transience of motor cortical activity at the onset of movement has several plausible explanations. The motor cortex may only be responsible for initiating movements at the right time in response to the learned cue^17^. It may also be necessary for generating a motor plan and, as movements are initiated, communicating it to other parts of the motor system that subsume control thereafter^18^. Finally, the motor cortex may be necessary for the first protrusion in a licking bout, when animals use their tongues to localize the reward port in space^16^, but not thereafter. In this interpretation, the trajectory of the tongue may be modified online if it does not immediately contact the target as intended, an interpretation supported by the observation that initial protrusions are often longer in duration (**Fig. 1b**), and more complex kinematically^16^ until the port is contacted. Then, once the port is localized, subsequent ballistic protrusions may be mediated by other motor centers.

To examine whether any of these hypotheses potentially explain the transience of cortical activity during bouts of tongue movements, we examined a separate cohort of animals trained to perform a ‘Delayed Reward Task’ that was identical to the Simple Reward Task, except that reward delivery was delayed until the 4^th^ port contact (C4) on 50% of trials (**Fig. 2a**). If the transience of motor cortical control in bouts of licking follows from a restricted role in (1) movement planning and initiation, or (2) port localization/trajectory modification, then we would expect to observe motor cortical disengagement with similar timing in the Simple and Delayed Reward Tasks. Examining behavior in this task, we found that, longer-duration, more variable and kinematically complex tongue movements persisted until reward delivery, well after the first port contact and after it is localized in space (**Fig. 2b**; **EDFig. 1**; **Supplementary Video 3**).

**Figure 2 -.**
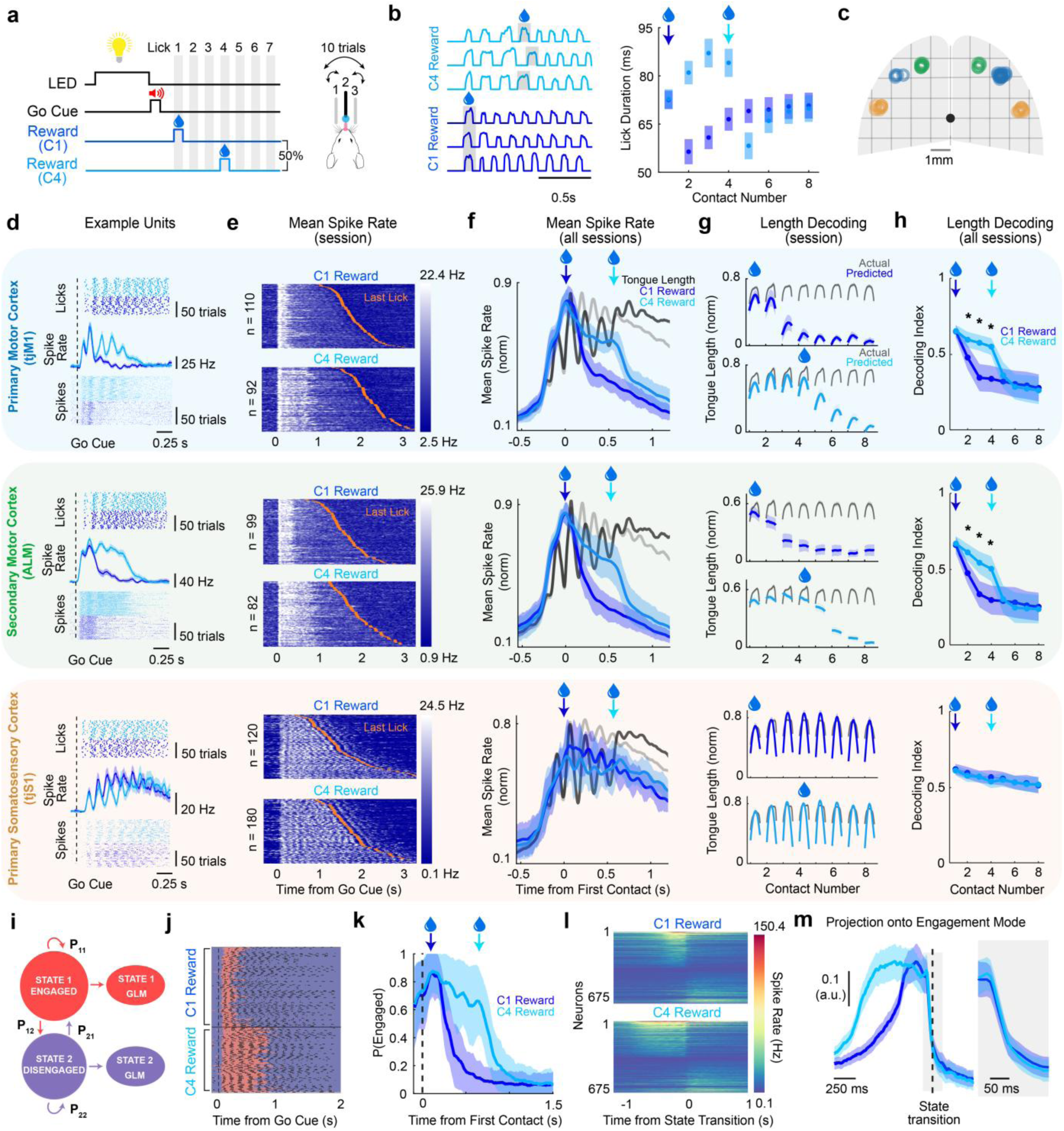
Motor cortical engagement in the Delayed Reward Task. **a.** Schematic of the Delayed Reward Task. **b.** Tongue length on exemplar trials (*left*) and the mean duration of the first eight tongue protrusions across sessions (*right; n = 36 sessions, 7 animals*). **c.** Location of all probe insertions targeted to the tjM1(*blue*), ALM (*green*), and tjS1 (*yellow*). **d.** Rasters of port contacts (*top*), spikes (*bottom*) and trial-averaged spike rate (*middle)* for example units in the tjM1 (*top row*), ALM (*middle row*), and tjS1 (*bottom row*) on each trial type. **e.** Mean spike rate across all units in the tjM1 (*top*), ALM (*middle*), and tjS1 (*bottom*) on C1 and C4 reward trials across an example session. Trials sorted by the timing of the last lick in a bout (*orange ticks*). **f.** Mean session-normalized spike rates on each trial type in the tjM1 (*top; n = 21 sessions, 6 animals*), ALM (*middle; n = 13 sessions, 4 animals*) and tjS1 (*bottom; n = 14 sessions, 3 animals*) across all sessions. Mean tongue length on C_1_ reward trials (*gray*) and C_4_ reward trials (*black*) indicated for reference. **g.** Tongue length predictions from the tjM1 (*top*), ALM (middle), and tjS1 (bottom). **h**. Decoding index (see ***Methods***) across all sessions (asterisks: p<0.005 comparing trial types, two-tailed t-test). **i.** Schematic of the HMM-GLM framework for inferring engaged and disengaged states. **j.** GLM-HMM state occupancy (*red*: engaged; *purple*: disengaged) in an example session. **k.** Posterior probability of the engaged state for C1 and C4 reward trials. **l.** Average spike rate of all units that have significantly different spike rates before and after the state transition (paired one-tailed t-tests for both increase and decrease). **m.** Projection of activity onto engagement mode (see *Methods*) for C1 and C4 trials across all sessions. Zoomed in (±100 𝑚𝑠 from transition) plot of the projection (inset). Shaded regions and error bars indicate 95% confidence intervals in all panels.

We again recorded activity in the tjM1 (n=934 single units, 21 sessions, 6 animals) and ALM (n=704 single units, 13 sessions, 3 animals) and found that motor cortical activation was also prolonged on C4-reward trials compared to C1-reward trials (**Fig. 2c**; **EDFig. 3**). In both cases, the timing of motor cortical disengagement was closely linked to the timing of reward delivery, which occurred well after both movement initiation and port localization/trajectory re-aiming on C4-reward trials. Prolonged activation on C4-reward trials was evident in the activity of single units (**Fig. 2d**) and at the level of population firing rates (**Fig. 2e,f**). Further, a decoder trained on the first tongue protrusion remained predictive of movement kinematics until reward delivery on C4-reward trials (**Fig. 2g,h**; **EDFig. 4**). In both trial types, motor cortical disengagement still occurred well before the cessation of movement. These observations indicate that the disengagement of the motor cortex from the control of movement does not appear to be a function of its transient involvement in movement initiation and/or port localization.

To assess whether cortical disengagement was a general feature of sensorimotor cortices, we also examined neural activity in the tongue/jaw primary somatosensory cortex (tjS1; n= 373 single units, 14 sessions, 3 animals; **Fig. 2c**). Units in the tjS1 were strongly activated during movement, but no signature of disengagement was observed at any point, regardless of reward timing. Rather, the activity of single units (**Fig. 2d**) and the overall population (**Fig. 2e,f**) was sustained until the cessation of movement. Tongue/jaw kinematics could also be decoded accurately for the duration of each bout of movement when a decoder was trained on the first tongue protrusion (**Fig. 2g,h**; **EDFig. 4**).

We evaluated whether state transitions on C1- and C4-reward trials, which occur at different times, were similar dynamically. To precisely identify the timing of state transitions on a trial-by-trial basis, we fit neural and kinematic data to a generalized linear model–hidden Markov model (GLM-HMM) that included two discrete states: one describing an engaged state of motor cortex and one describing a disengaged state of motor cortex (**Fig. 2i**). Here, each discrete state defines a state-specific Gaussian GLM encoding model that relates kinematics to population neural activity. We found that the engaged state was occupied for a prolonged duration when water was delivered following the 4^th^ port contact (C4) versus the 1^st^ port contact (C1; **Fig. 2j,k**; **EDFig. 6**). Examining the activity of single units relative to the time of cortical disengagement identified by the HMM-GLM revealed the presence of a discrete state switch in neural dynamics that was highly similar across C1- and C4-reward trials but occurred at different times relative to movement onset (**Fig. 2l**). To determine the timescale over which disengagement occurred, we computed an ‘engagement mode’ of neural activity – the linear combination of neural activity with the largest change in activity between the engaged and disengaged states (comparing the 400 ms prior to vs. the 400 ms after the state transition on C4-reward trials) – and projected neural activity onto this mode. The transition between the engaged and disengaged states occurred abruptly (90-10% decay time: 85 ± 23 ms; n = 35 sessions; 6 animals). No difference was observed in the magnitude or duration of decay in activity along this mode across trial types (**Fig. 2m**).

### Persistent motor cortical engagement during reward consumption

An alternative hypothesis that could explain why the motor cortex disengages from the control of movement is that the availability of water induces a switch to a cortically independent motor program responsible for the consumption of water, an innate behavior^9,11,19,20^. This hypothesis predicts that motor cortical disengagement is triggered by water delivery, consistent with the observation that the timing of disengagement is delayed when the timing of reward is delayed.

To test this hypothesis, we trained a new cohort of animals on a ‘Double Reward Task’ (**Fig. 3a**) and then recorded neural activity from the ALM and tjM1 (**Fig. 3c**, **EDFig. 3**) as mice performed the task. This task again followed a similar structure, but with another modified reward schedule. In this task, animals always received a reward at the first port contact (C1) but also received a second reward following the sixth port contact (C6) on 50% of trials (C1 and C1,6-reward trials; **Fig. 3a**). The duration and kinematic complexity of tongue protrusions reflected the altered task structure. The duration of protrusions decreased after the first reward delivery when water was being consumed, as in other tasks (one-tailed t-test, p<0.005, **Fig. 3b**; **EDFig. 1**), before increasing again prior to the second reward (one-tailed t-test, p<0.005).

**Figure 3 -.**
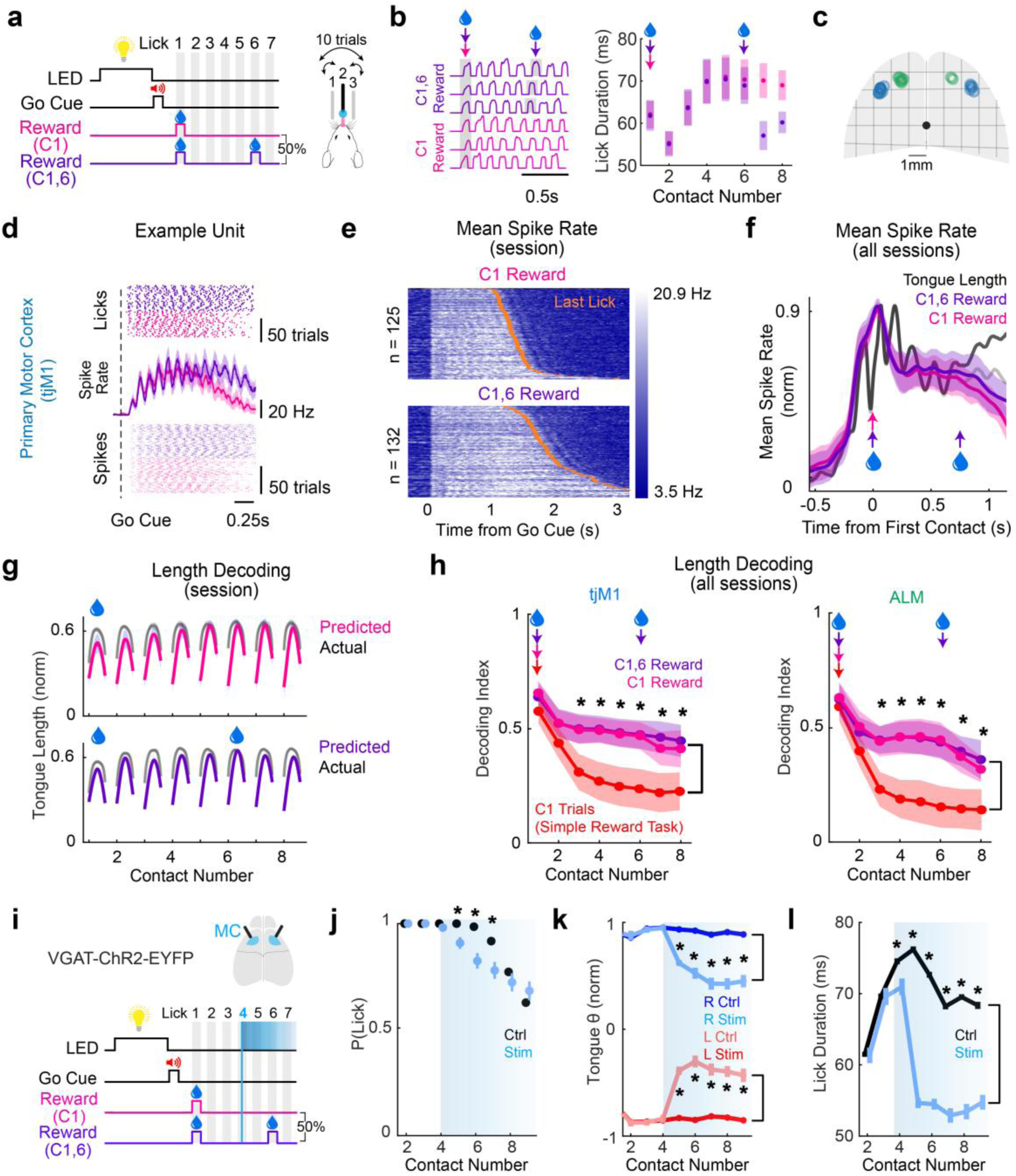
Motor cortical engagement in the Double Reward Task. **a.** Schematic of the Double Reward Task. **b.** Tongue length on exemplar trials (*left*) and the mean duration of the first eight tongue protrusions across sessions (*right; n = 26 sessions, 6 animals*). **c.** Location of all probe insertions targeted to the tjM1(*blue*) and ALM (*green*). **d.** Rasters of port contacts (*top*), spikes (*bottom*) and trial-averaged spike rate (*middle)* for example units in the tjM1 (*top row*), ALM (*middle row*), and tjS1 (*bottom row*) on each trial type. **e.** Mean spike rate across all units in the tjM1 (*top*), ALM (*middle*), and tjS1 (*bottom*) in an example session. Trials sorted by the timing of the last lick in a bout (*orange ticks*). **f.** Mean session-normalized spike rates on each trial type in the tjM1 across all sessions (*n = 18 sessions, 4 animals)*. Mean tongue length on C1 reward trials (*gray*) and C1,6 reward trials (*black*) indicated for reference. **g.** Tongue length predictions from the tjM1 in an example session. **h.** Decoding index (see ***Methods***) across all sessions (asterisks: p<0.005 comparing C_1_ trials in the Simple Reward Task and Double Reward Task; two-tailed t-test) for the tjM1 and ALM (*n = 15 sessions, 4 animals)*. **i.** Schematic of tjM1/ALM photoinhibition in the Double Reward Task. **j.** Probability of tongue protrusion on control (*black*) and photoinhibition trials (*light blue*; asterisks: p<0.005, two-tailed t-test, *n = 16 sessions, 4 animals*). **k.** Mean tongue angle on left (*red*) and right (*blue*) trials. Lighter shades represent photoinhibition trials (asterisks: p < 0.005 comparing photoinhibition trials to control trials; two-tailed t-test). **l.** Mean duration of tongue protrusions on control and photoinhibition trials (asterisks: p < 0.005, two-tailed t-test). Shaded regions indicate 95% confidence intervals. Shaded regions and error bars indicate 95% confidence intervals in all panels.

To determine whether motor cortical disengagement also followed the first port contact during the consumption of water, we again examined neural activity in the tjM1 (n=1213 single units, 18 sessions, 4 animals) and ALM (n=813 single units, 15 sessions, 5 animals). Surprisingly, the introduction of the variable second reward caused the motor cortex to persist in the engaged state, in both the tjM1 and ALM. The absence of disengagement was apparent at the level of single units (**Fig. 3d**; **EDFig. 7**), the population firing rate (**Fig. 3e,f**; **EDFig. 7**), and in decoding analyses (**Fig. 3g,h**; **EDFig. 7**). Not only was no disengagement observed following consumption of the first reward, but motor cortical engagement typically persisted past the second reward until the termination of movement. Even though we observed a transition to shorter-duration consumptive tongue movements whenever water was delivered, as in other tasks, this change in action was not accompanied by a transition to the disengaged state.

We next sought to determine whether this persistent motor cortical engagement in the Double Reward Task reflected a prolonged period of active cortical control of movement. We re-trained the same cohort of mice in which photoinhibition was performed in the Simple Reward Task. This re-training involved only the introduction of a second reward in response to the 6^th^ port contact on 50% of trials (**EDFig. 2**). Photoinhibition (10% of trials) was again applied on the 4^th^ lick contact. Now, following this additional period of training, silencing motor cortical activity reduced the rate of tongue protrusions (**Fig. 3j**; n = 4 animals, 18 sessions). When the tongue did protrude, animals again could not direct movements laterally to reward ports (**Fig. 3k**; n = 4 animals, 18 sessions; **Supplementary Video 4**). Here, C1-reward trials proceeded in an identical fashion in the Simple Reward Task and Double Reward Task, yet the effect of photoinhibition differed dramatically – in the same animals – across these contexts. These trials only differed in whether a second reward was possible. Only when there was a possibility of future reward did photoinhibition impair movement. These results suggest that motor cortical disengagement is unlikely to reflect a switch to an innate consummatory motor program, as consumption proceeded without any observed disengagement.

### Dissociating motor cortical disengagement from reward

In each task, the motor cortex remained engaged in controlling movements so long as animals expected that a water reward might be delivered. One explanation for this observation is that motor cortical engagement is associated with a motivated state or reward prediction signal that persists so long as future rewards are anticipated. That is, engagement may follow from anticipation of water delivery, a positive valence event, and not from a process explicitly related to the control of movement.

Notably, however, animals must switch between two related motor programs at the time of reward in paradigms in which animals lick for water rewards. First, the tongue is used to detect the presence of reward. Then, once water is sensed on the tongue, it must be brought to the mouth and swallowed. Disengagement does not occur during reward consumption in the Double Reward Task suggesting that motor cortical disengagement is not a direct consequence of this switch between programs. Instead, the motor cortex may remain engaged so long as animals anticipate a sensory-guided switch in motor programs (exploratory palpation to consummatory lapping) that is contingent upon a sensory event (the sensation of water on the tongue). In this latter interpretation, motor cortical engagement is explicitly tied to this sensorimotor process and could indicate a state of ‘sensorimotor attention’ that persists so long as animals are actively awaiting a sensory event – whose delivery and timing is uncertain – upon which a future switch in motor action is contingent. Once future rewards are no longer anticipated, animals need not anticipate a sensation-contingent switch in actions and therefore may not need to attend to the sensorimotor process.

To determine whether disengagement is explicitly related to the expectation of future reward, or an expectation of an upcoming switch in motor programs, we trained another cohort of animals on a ‘VTA Reward Task’ that explicitly decouples reward from associated changes in movement. Here, animals were rewarded only with optogenetic stimulation of dopaminergic neurons in the ventral tegmental area (VTA), rather than with a water reward that is inherently linked to a switch to consummatory movements (**Fig. 4a,b**). In the VTA Reward Task, reward (bilateral VTA stimulation) was initiated in response to licking and was delivered on the fourth port contact (C4; **Fig. 4a,b**). Animals licked for VTA stimulation alone and ceased to lick within a few trials if VTA stimulation was absent.

**Figure 4 -.**
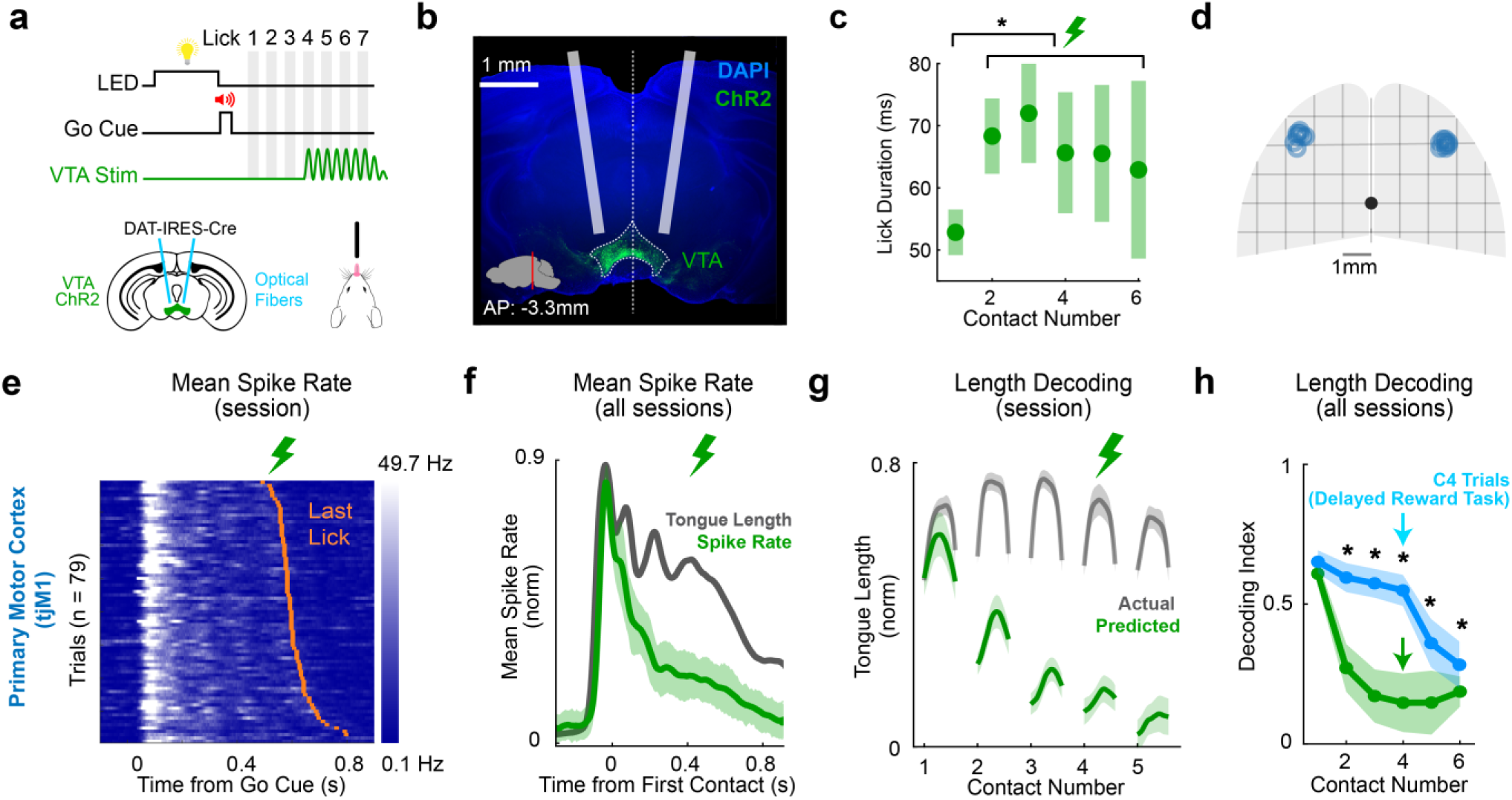
Motor cortical engagement in the VTA Stimulation Task. **a.** Schematic of the VTA Stimulation Task (*left*) and experimental approach used for optogenetic activation of VTA dopaminergic neurons (*right*). **b.** Example histological image illustrating optic fiber placement and viral expression of ChR2-EYFP (*green*) in the VTA. **c**. Mean duration of the first six tongue protrusions across sessions (*n = 18 sessions, 4 animals)*. The duration of each subsequent tongue protrusion was increased relative to the first (p<0.005; two-tailed t-tests comparing each protrusion to the first). **d.** Location of all probe insertions targeted to the tjM1(*blue*). **e.** Mean spike rate across all units in the tjM1 in an example session. Trials sorted by the timing of the last lick in a bout (*orange ticks*). **f.** Mean session-normalized tjM1 spike rate (*green*) and average tongue length (*black*) across all sessions (*n = 18 sessions, 4 animals)*. **g.** Tongue length prediction from tjM1 recordings in an example session. **h.** Decoding index (normalized model RMSE, see ***Methods***) across all sessions (asterisks: p<0.005 comparing C_4_ trials in the VTA Stimulation Task (*green*) and Delayed Reward Task (*blue*); one-tailed t-test). Shaded regions indicate 95% confidence intervals. Shaded regions and error bars indicate 95% confidence intervals in all panels.

We recorded neural activity in the tjM1 (n = 719 single units, 18 sessions, 4 animals) to determine if disengagement remained linked to reward timing in this task (**Fig. 4d**). We found that activity increased transiently at movement onset, as in other tasks, but rapidly decayed after the first protrusion, well before the time of reward delivery (**Fig. 4e,f**). Tongue/jaw kinematics could be predicted accurately only during the first tongue protrusion (**Fig. 4g,h**). The fast decay of activity immediately following movement onset in this paradigm indicates that motor cortical engagement is not inherently linked to motivation and/or the anticipation of the positive valence event. Instead, motor cortical engagement only persisted when a reward that necessitated a switch in action was anticipated. In the VTA Reward Task, when there was no expectation of an upcoming sensation-contingent switch in motor program, motor cortical activation only accompanied the onset of movement.

Notably, in the VTA Reward Task, the duration and complexity of protrusions did not decrease after the port was localized in space, but rather increased following the first port contact, presumably reflecting a lack of consummatory movements (**Fig. 4c**; **EDFig. 1**). Thus, long-duration ‘exploratory’ movements and shorter-duration consummatory movements can both be executed in the engaged state (**Fig. 3b**) and can both be executed in the disengaged state (**Fig. 2b**; **Fig. 4c**).

### Motor cortical disengagement is shaped during task acquisition

We hypothesized that motor cortical engagement is a signature of a state in which animals are attentive to their movement. We further posit that, here, this attention is evoked by uncertainty – uncertainty about future actions that follows from uncertainty about the external world. In tasks motivated by water reward, uncertainty about future movements largely represents uncertainty about when and whether water will be delivered and sensed on the tongue. When animals anticipate that a tongue protrusion *could* touch water, they must be attentive to their actions and associated sensory outcomes. This interpretation yields two predictions about motor cortical engagement during task acquisition. First, engagement should be persistent during initial task exposure, when the delivery of water cannot be anticipated. Before animals learn the task structure, there is a high degree of uncertainty about what they are likely to feel and about the actions they must execute in the near future. Second, as animals learn when rewards will *not* be delivered, engagement should diminish in a commensurate fashion.

To test these predictions, we recorded activity in the tjM1 of an additional cohort of animals performing the Simple Reward Task over their first five consecutive days of task exposure (**Fig. 5a,b**, **EDFig. 3**; n = 41 +/- 14 single/units per session, 5 sessions/animal, 4 animals), a period during which trial-by-trial variability in kinematics decreased and approached that observed in expert animals (**EDFig. 1**). Once animals learn that water is only available upon the first port contact in this task, there should be little uncertainty about whether a reward will be sensed subsequently and no need to resolve uncertainty with tactile feedback to guide upcoming movements. Indeed, we found a high degree of cortical activation that persisted throughout the duration of movement on the first day of task exposure (**Fig. 5c-e**). Further, as predicted, we observed a marked reduction in both motor cortical activity and in the degree to which kinematics could be decoded from neural activity following reward delivery across the first five days of training (**Fig. 5c-g**). By the fifth day of training, motor cortical engagement was highly transient, as we initially observed in expert animals performing the Simple Reward Task. The progressive emergence of a disengaged state during task acquisition further supports the interpretation that motor cortical disengagement follows when there is high confidence that no reward will be delivered – when future sensation and action are predictable and there is longer utility in attending to the relevant sensorimotor process.

**Figure 5 -.**
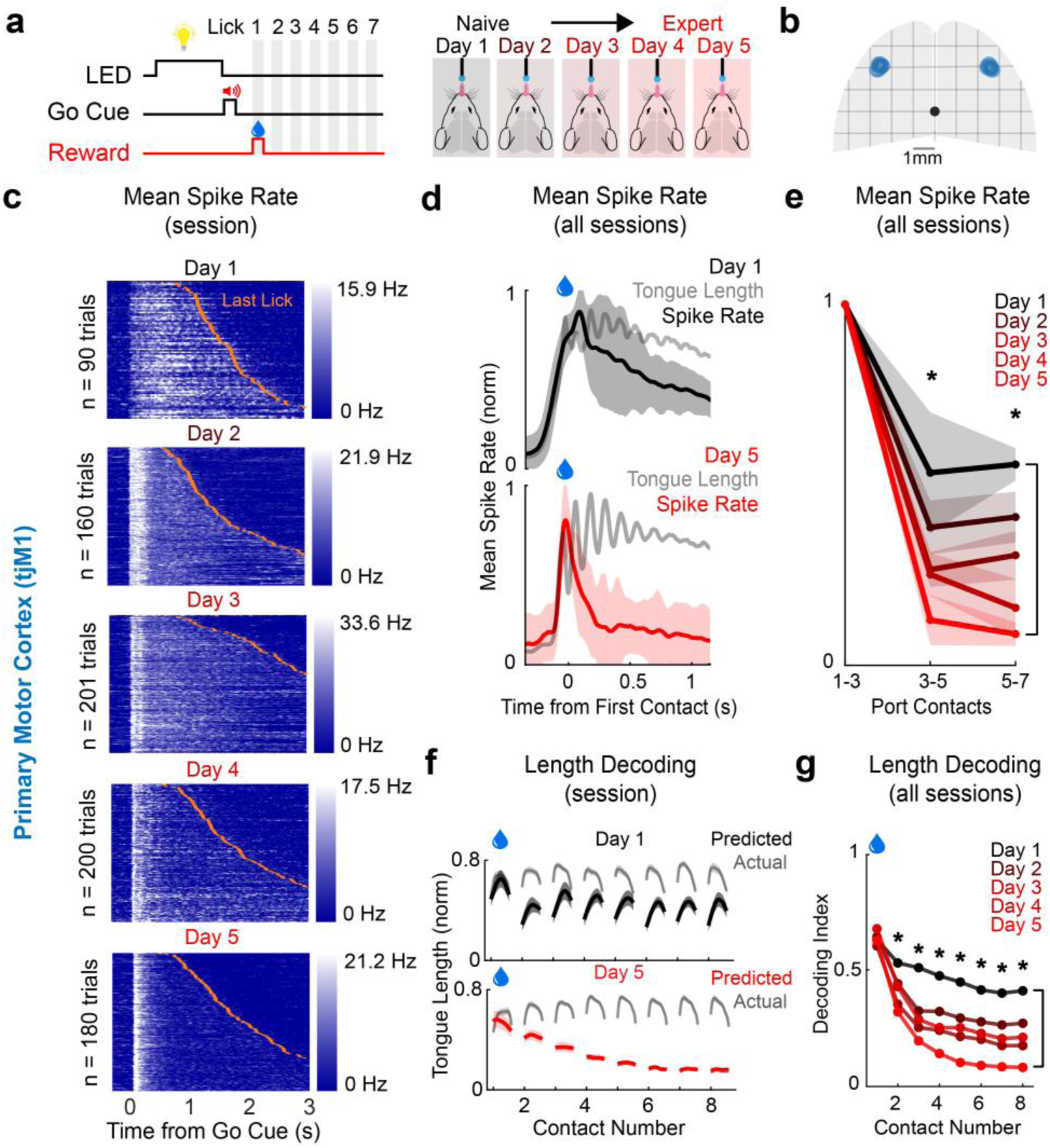
Motor cortical engagement during task acquisition. **a.** Schematic of experimental paradigm. Mice were first exposed to the Simple Reward Task on Day 1 and were observed across the first five consecutive days of task exposure. **b.** Location of all probe insertions targeted to the tjM1(*blue*). **c.** Baseline (up to 0.5 s prior to Go Cue) subtracted mean spike rate across all units in the tjM1 on example sessions on Days 1 - 5. Trials sorted by the timing of the last lick in a bout (*orange ticks*). **d.** Mean session-normalized spike rates in the tjM1 across all Day 1 (*top; n = 4 sessions, 4 animals*) and Day 5 (*bottom; n = 4 sessions, 4 animals*) sessions. **e.** Mean session-normalized spike rate on each day of training, binned across protrusions (to account for the irregularity of movement on some Day 1 sessions). Asterisks indicate a significant reduction in spike rate between Day 1 and Day 5 (p < 0.005, one-tailed t-test). **f.** Tongue length prediction from tjM1 recordings in example Day 1 (*top*) and Day 5 (*bottom*) sessions. **g.** Decoding index (normalized model RMSE, see ***Methods***) across all sessions for each day of task exposure (asterisks: p<0.005 comparing Day 1 and Day 5, one-tailed t-test). Shaded regions and error bars indicate 95% confidence intervals in all panels.

## DISCUSSION

The necessity of the motor cortex for movements can vary across actions and across learning^9,10,13–15,21–23^. We demonstrate that the cortical control of movements can also vary on a faster timescale, engaging and disengaging discretely on sub-second timescales even during ongoing movements that otherwise appear continuous. We posit that whether and when these transitions occur is a function of animals’ uncertainty about upcoming sensorimotor processes.

To examine transitions in the cortical control of movements, we examined a set of head-fixed licking tasks. In a task in which water reward was always delivered immediately upon contact with a reward spout, the motor cortex was engaged only transiently, with motor cortical activity – and the influence of motor cortical activity on movement – decaying rapidly after movements were initiated (**Fig. 1**). However, when the timing of the single reward was uncertain (**Fig. 2**), motor cortical engagement was sustained until the reward was received, suggesting that the transience of motor cortical activation was not simply tied to the initiation of movement or to the initial localization of the reward port in space. When the number of rewards was varied, so that there was a persistent possibility of reward, the motor cortex remained persistently engaged as well, typically until the cessation of movement (**Fig. 3**). In this context, cortical activation persisted even while water was being consumed. Thus, water consumption – an innate, highly practiced action – occurred with the motor cortex highly engaged when this action was executed in a context marked by uncertainty about future events. Importantly, engagement of the motor cortex was not linked to reward explicitly. When mice received an internal reward – the activation of dopaminergic neurons in the VTA – motor cortical disengagement decoupled from the timing of reward (**Fig. 4**). This suggests that cortical engagement persisted until reward not because of an expectation of a positive-valence event, but because of anticipation of a switch in motor programs triggered by the sensation of reward. The interpretation that cortical engagement in movement is induced by uncertainty was supported by observations during learning. The disengaged cortical state emerged over time as uncertainty about the reward schedule decreased (**Fig. 5**).

Across all tasks, we found that the motor cortex was reliably disengaged from movement when – and only when – there was no further expectation that a sensory guided switch in motor actions would be necessary in the immediate future. A parsimonious explanation for these results is that motor cortical engagement reflects a state of sensorimotor attention, or ‘attention to movement,’ that follows when future actions and/or their sensory consequences are unpredictable. In behavioral contexts in which actions are contingent upon uncertain environmental features or events, animals must direct attention to movement processes and their associated sensory consequences. For example, when one traverses an icy sidewalk, where a loss of traction may occur in an unpredictable fashion, close attention must be paid to that sensorimotor process. Tactile and proprioceptive feedback – the sensory consequences of the movement – must be monitored so that slips can be quickly identified and appropriate actions engaged to regain balance. In the tasks examined here, uncertainty principally takes the form of uncertainty about when rewards will be delivered – not because of the positive valence of the reward itself, but because the sensation of water prompts a switch in the appropriate motor program – a switch from an exploratory action in which mice palpate the port to monitor if and when water is present to a consummatory program in which water must be lapped to the mouth and swallowed. Uncertainty may also impact motor planning. When there is uncertainty about future actions, multiple competing motor plans may be encoded until the most appropriate action is selected and initiated. However, the direct effect of motor cortex photoinhibition on the kinematics of ongoing movements suggests this is unlikely the sole explanation for motor cortical engagement.

Notably, motor cortical engagement and disengagement was not tied to any detectable change in how sensory information was represented in the primary somatosensory cortex, where activity faithfully reflected movements in an invariant manner. Thus, while the semantic concept of attention appears germane to the phenomenology of cortical disengagement reported here, these observations suggest the variable activation of the motor cortex during movement is independent from the oft-studied modulation of sensory representations^24–28^. We also note that movement initiation was always contingent upon perception of an auditory go cue that itself was preceded by the illumination of a ‘ready’ LED. One might therefore expect auditory attention prior to movement, but motor cortical activation was not observed prior to movement onset. Thus, a generalized attentional state does not appear explanatory of motor cortical activation per se.

Another notable feature of our results was the persistent activation of the motor cortex beyond the second, and final, reward in the Double Reward Task. In this task, there is zero probability of an additional reward once the second reward is delivered. Thus, interpreting motor cortical engagement as reflecting sensorimotor uncertainty would predict disengagement following the second reward delivery. The persistent engagement we observed in this task was unexpected but could indicate that mice did not track the cumulative number of rewards delivered and update their expectations accordingly.

Tongue movements in rodents are controlled by one or more central pattern generators (CPGs)^29–32^. CPG-mediated movements – which include breathing, chewing, drinking, suckling, swallowing, vocalizing, locomoting and copulating – represent a large class of movements that are most critical for all mammals to survive and thrive^30,33–37^. Nevertheless, the control of CPG-associated movements differs from other movements, and from movements in other mammals, to varying degrees. Whether the motor cortex dynamically engages and disengages during other movements in rodents, and/or during movements in other mammals remains an open question. Some evidence suggests an analogous process during locomotion in cats and mice – where motor cortical activity emerges selectively when animals encounter obstacles, a context in which motor cortical lesions also selectively impair behavior^21,22,38^. Functional neuroimaging has demonstrated that attention to movements and/or targets can modulate the activation of sensorimotor cortices in humans^39–43^. Thus, attention may be a critical determinant of the neural systems invoked to control movement. Future work examining how movements are controlled across different neural states and how state switches are instantiated across the motor system should prove highly insightful.

## ACKNOWLEDGMENTS

The authors thank Jaclyn Birnbaum and Daniel O’Connor for helpful discussions. We thank Jaclyn Birnbaum, Nuo Li, Hidehiko Inagaki, Tim Wang, and Yujin Han for helpful comments on the manuscript. This work was supported by the Whitehall Foundation, the Klingenstein Fund, the Simons Foundation, NIH R01NS121409, NIH U19NS137920, and NSF 2239412.

## AUTHOR CONTRIBUTIONS

TD and MNE conceived the project. MNE and BD supervised research. TD and MNE designed experiments. TD performed experiments. TD, ZFL, and MAH analyzed data. MAH provided technical assistance. TD, ZFL, and MNE wrote the manuscript with input from all authors.

## COMPETING INTERESTS

The authors declare no competing interests.

## MATERIALS AND METHODS

### Animals

This study used data collected from 27 mice; both male and female animals between 8 weeks and 13 weeks of age were used. Four animals were used for the Simple Reward task, seven animals were used for the Delayed Reward task, six animals were used for the Double Reward task, and four animals were used for the Simple Reward task Learning experiments and were either C57BL/6J (The Jackson Laboratory (JAX), 000664) or VGAT-ChR2-EYFP (−/−) (bred in-house; breeders: JAX, 014548). Four VGAT-ChR2-EYFP (+/−) animals were used in photoinhibition experiments (two animals were used in electrophysiology recordings in the Double Reward task). Finally, four DAT-IRES-Cre (The Jackson Laboratory (JAX), 006660) (+/+) animals were used in the VTA stimulation task experiments. Mice were housed in a 12 h reverse dark/light cycle room (ambient temperature of 68–79 °F (20–26 °C); humidity of 30–70%) with ad libitum access to food. Access to water was restricted during behavioral and electrophysiology experiments (see ‘Mouse Behavior’ subsection).

### Surgical procedures

All surgical procedures were performed in accordance with protocols approved by the Boston University Institutional Animal Care and Use Committee. For postoperative analgesia, mice were given ketoprofen (0.1 ml of 1 mg ml^−1^ solution) and buprenorphine (0.06 ml of 0.03 mg ml^−1^ solution) before the start of all surgical procedures. Mice were anesthetized with 1–2% isoflurane and placed on a heating pad in a stereotaxic apparatus. Artificial tears (sodium chloride 5% opthalmic ointment; Akorn) were applied to their eyes, and a local anesthetic was injected under the skin (bupivacaine; 0.1 ml of 5 mg ml^−1^ solution) above the skull. The skin overlying the skull was removed to expose the ALM (AP: +2.5 mm, ML: ±1.5 mm), tjM1(AP: +2.2 mm, ML: ±2.5 mm), and/or tjS1 (AP: +0.5 mm, ML: ±3.5 mm), bregma and lambda. The periosteum was removed and the skin was secured to the skull with cyanoacrylate (Krazy Glue) around the margins. For electrophysiology experiments, a headbar was implanted just anterior to bregma and secured with superglue and dental cement (Jet). Wells to hold cortex buffer (NaCl 125 mM, KCl 5 mM, glucose 10 mM, HEPES 10 mM, CaCl_2_ 2 mM, MgSO_4_ 2 mM, pH 7.4) during electrophysiology recordings were sculpted using dental cement, and a thin layer of superglue was applied over any exposed bone.

For VTA stimulation experiments, virus (AAV9-EF1α-DIO-hChR2(H134R)-EYFP; Addgene plasmid #20298; titer 1.3×10^13^ vg/mL) was injected into the VTA using a custom-made manual volume displacement injector connected to a glass pipette (MO-10; Narishige) pulled to a 30 µm tip (P-2000; Sutter Instruments) that was beveled to a sharp tip. Pipettes were back-filled with mineral oil and virus was front-loaded before injection. Pipettes were inserted through the thinned bone to the appropriate depth and virus injected at 10 nl min^−1^. 150nL of virus was injected bilaterally into the VTA (AP: -3.3 mm, ML = ±0.9 mm, DV = +4.4 mm) at a 9^°^ angle on the medial-lateral axis of the head leveled in the anterior-posterior axis. Fiber optic cannulas (MFC_200/470-0.37_5.0mm; Doric Lenses) were implanted 0.15 mm over the target region and a headbar was implanted anterior to Bregma. Dental acrylic (Jet repair; Pearson Dental) was used to secure the optic fiber and headbar to the skull and protect exposed bone.

For photoinhibition experiments, after headbar implantation, bone overlying ALM and tjM1 (AP: +2.15 mm, ML: 2 mm, radius = 0.5 mm) was thinned bilaterally with a dental drill and a layer of cyanoacrylate adhesive and Metabond was thinly layered over the skull and left to dry. Fiber optic cannulas (MFC_400/470-0.37_1.0mm; Doric Lenses) were superficially implanted over the target region and a headbar was implanted anterior to Bregma. Dental acrylic (Jet repair; Pearson Dental) was used to secure the optic fiber and headbar to the skull and protect exposed bone.

### Histology

Mice were perfused transcardially with phosphate-buffered saline (PBS) followed by 4% PFA in 0.1 M PB. The tissue was fixed in 4% PFA at least overnight. The brain was then suspended in 3% agarose in PBS. A vibratome (Compresstome VF-510-0Z; Precisionary) cut coronal sections of 100 μm that were mounted and subsequently imaged on a fluorescence microscope (BX41, Olympus). Images showing CM-DiI fluorescence (C7001; ThermoFisher) were collected to recover the location of silicon probe recordings. Images showing GFP expression were collected to confirm the targeting of virus injection and optic fiber implantation over the VTA.

### Mouse behavior

After surgery, mice were allowed to recover for approximately 1 week and then water restricted, receiving 1.0 mL of water per day. Animals used for the VTA stimulation task were given approximately 1.0 mL of water. Behavioral training commenced after animals had been water restricted for 3–5 days. If animals received less than 1.0 mL during training, they were supplemented with additional water.

The behavioral system was by controlled by a state machine running on a microcontroller (Bpod r0.5; Sanworks) defined using custom scripts in MATLAB. Tongue contacts with the port were detected using a two-transistor conductive lick detector(https://www.janelia.org/open-science/dual-lick-port-detector)^44^ which registered a measurable voltage change when the mouse completed the circuit by touching the port.

For all tasks, animals were headfixed in a tube and trained to lick a port to receive a reward. Each trial began with the onset of a 2 s LED cue. Following the offset of the LED, an auditory Go Cue in the form of a 10 ms swept-frequency cosine (chirp) sound was played, signaling that the animal could lick the port to obtain a reward. If the animal contacted the lickport during the LED period, the current trial epoch was reset, ensuring that they withheld licking for the full 2 s before proceeding to the next stage.

In the Simple Reward task, animals were rewarded on the first port contact. In the Delayed Reward task, rewards were delivered on either the first or fourth port contact with equal probability (50%). In the Double Reward task, animals received a reward either on the first contact alone or on both the first and sixth contacts, also with 50% probability. In the VTA Stimulation task, reward was induced optically on the fourth port contact. For all tasks except the VTA Stimulation task, the reward consisted of approximately 2 µl of water. In the VTA Stimulation task, the water reward was replaced by 0.9 s optogenetic stimulation of dopaminergic neurons in the ventral tegmental area (VTA). In the Simple Reward and Delayed Reward Tasks, the lickport was moved amongst three lateral positions in a block-wise manner, switching every 10 trials. In some animals performing the Double Reward Task (n=4; all used for electrophysiology), the lickport similarly switched between three lateral positions every 10 trials. In other animals (n=2; used only for photoinhibition; n=2 used for photoinhibition and electrophysiology), the lickport switched between two lateral positions (omitting the central position) every 20 trials. In the learning paradigm, the lick port was held in a fixed, centrally located position, as this was the first stage of training for all animals in all tasks. In the VTA stimulation task, the port was also located in a single central position for simplicity.

Two perpendicular linear stages (LSM050B-T4 and LSM025B-T4; Zaber) provided motorized positioning of the lickport along the anteroposterior and mediolateral axes. Vertical adjustment was achieved with a manual linear stage (MT1/M; Thorlabs). Both stages were driven by an X-MCB2 controller (Zaber), which received movement commands over a serial link from a microcontroller (Teensy 3.2; PJRC). Port locations were defined relative to a chosen origin and arranged along an arc centered on the midline, with equal spacing in arc length between adjacent positions.

For all tasks, animals were first habituated to the head fixation apparatus for 15 minutes prior to the start of training. They initially learned to associate licking with reward by receiving water coincident with the presentation of a high-pitched auditory tone following the LED cue. After approximately 20 trials, water was no longer automatically delivered but instead was triggered by a lick port contact following the tone. After three days of training, the lick port began shifting between all positions in block wise fashion. The distance between port positions was gradually increased over time. Following this, the same set of port positions was held fixed for an additional one to two weeks of training to ensure expert level performance.

In the Delayed Reward and Double Reward tasks, modified reward timing was introduced by randomly interleaving trial types over two days, with the port fixed directly in front of the animal. After this period, the port began switching between lateral positions in blocks. For the VTA stimulation task, animals initially received water paired with the high-pitched auditory Go Cue tone on the first day of training for 10 to 50 trials, during which VTA stimulation was also delivered on the first lickport contact. After this initial phase, water was no longer automatically delivered but was triggered by the first port contact following the tone. Over the next 50 to 100 trials, the water reward was gradually reduced to zero, after which animals consistently licked in response to VTA stimulation alone. Over the course of 3 to 10 weeks, the timing of VTA stimulation was progressively delayed from the first to the fourth port contact. Animals were considered expert once they had completed one week of training with VTA stimulation occurring on the fourth contact.

Mice performed the task in darkness with no visual cues relating to the position of the port. To prevent mice from using sounds emitted by the motor to guide their behavior, we played a constant white noise (cut-off at 40 kHz; 80 dB SPL). The constant white noise was also played in tasks in which the port was not moved to ensure consistency across tasks.

### Videography analysis

High-speed video was captured (400-Hz frame rate) from two cameras (FLIR, Blackfly monochrome camera, BFS-U3-16S2M-CS). One provided a side view of the mouse, and the other provided a bottom view. We tracked the movements of specific body features using DeepLabCut^45^. The tongue, jaw and nose were tracked using both cameras. Paws were tracked using the bottom view. Position and velocity of each tracked feature was calculated from each camera. The *x* and *y* position of each kinematic feature was extracted from the output of DeepLabCut. Missing values were filled in with the nearest available value for all features, except for the tongue.

### Photoinhibition experiments

Optogenetic photoinhibition was performed on approximately 10% of trials selected at random. A ‘masking flash’ (470-mm LEDs; LUXEON Rebel LED) controlled by a microcontroller (Nano 33 BLE Sense; Arduino) was delivered continuously (100 ms, 10 Hz) for the duration of the session to prevent mice from differentiating control and photoinhibition trials.

Photoinhibition was performed through a clear skull approach (see ***Surgical procedures***). Bilateral stimulation of the brain was achieved using a pair of optic fibers (0.39 NA, 400 µm core diameter) that were implanted above the clear skull during surgery. These optic fibers were coupled to 470 nm LEDs (M470F3; Thorlabs) which were attached before the beginning of each behavioral session. Light was delivered at the onset of the fourth port contact for 0.7 s, followed by a 0.2 s linear ramp down. We targeted ALM and tjM1 bilaterally by stimulating using a sinusoidal wave with a frequency of 40 Hz. The total power of light delivered through the optic fibers was 8 mW.

### VTA Stimulation

Optogenetic photoactivation of neurons expressing the DAT protein in the VTA was delivered on 100% of trials as reward in lieu of water reward. A ‘masking flash’ (470-mm LEDs; LUXEON Rebel LED) controlled by a microcontroller (Nano 33 BLE Sense; Arduino) was delivered continuously (100 ms, 10 Hz) for the duration of the session. Black tape was used to prevent light leakage from the optic fiber. Bilateral stimulation of the brain was achieved using a pair of optic fibers (0.37 NA, 200 µm core diameter) that were implanted above the VTA during surgery. These optic fibers were coupled to 470 nm LEDs (Thorlabs; M470F3) which were attached before the beginning of each behavioral session using a ceramic coupler. Instead of water reward, light was delivered at the onset of the fourth port contact for 0.7 s (6 mW per fiber, 40 Hz sinusoid), followed by a 0.2 s linear ramp in amplitude.

### Behavioral analysis

Behavioral data were collected using the Bpod (r0.5; Sanworks). All sessions used for behavioral analysis had at least 140 total completed trials for water reward tasks, 70 trials for learning experiments, and 60 trials for VTA stimulation experiments.

Tongue angle and length were found using the bottom camera view. Tongue angle was defined as the angle between the vector pointing from the jaw to the tip of the tongue and the vector defining the direction the mouse was facing. Tongue length was calculated as the Euclidean distance from the jaw to the tip of the tongue. All kinematic features were first standardized by taking the 99th percentile across time and trials and normalizing to this value. To quantify the duration of individual licks within a bout, we measured the total time the tongue was visible on a lick-by-lick basis throughout each trial.

Summary plots for lick duration were computed by determining, for each animal and session, the mean duration of each lick across trials. Session means were then averaged across sessions and animals to yield the group-level mean duration per lick, which was plotted with its 95% confidence interval, computed as ±1.96 times the standard error of the mean. Lick-to-lick variability was quantified similarly. For each animal and session, the standard deviation of the duration of each lick (i.e. 1^st^ lick, 2^nd^ lick, 3^rd^ lick, etc.) across trials was calculated, session-wise standard deviations were averaged across sessions and animals, and the resulting group-level mean standard deviation per lick was plotted with its 95% confidence interval, calculated in the same way.

### Electrophysiology recordings

Extracellular recordings were performed in cortical areas using Neuropixels 1.0 and 2.0 (1 shank), allowing for recordings from 384 channels arranged in a checkerboard pattern (IMEC). The signals were demultiplexed into 384 voltage traces sampled at 30 kHz and stored for offline analysis.

At least 6 h before recording, a small craniotomy (1–1.5 mm diameter) was made over one or more of the ALM (AP: +2.5 mm, ML: ±1.5 mm), tjM1(AP: +2.3 mm, ML: ±2.5 mm), and/or tjS1 (AP: +0.5 mm, ML: ±3.5 mm).

Then, 2–8 recordings were performed through a single craniotomy on consecutive days. After inserting the probes to a depth of 900 - 1,100 µm (ALM) or 1,200 - 1,400 µm for tjM1 and tjS1 (MPM System; New Scale Technologies), brain tissue was allowed to settle for at least 5 min before starting recordings. All recordings were made using SpikeGLX (phase3a release; https://billkarsh.github.io/SpikeGLX/).

AP-band signals were high-pass filtered at 300 Hz (third-order zero-phase Butterworth filter), and the local common mode signal - computed as the median across ±12 adjacent channels - was subtracted from each channel. Samples exceeding a high-power threshold were blanked to suppress artifacts.

### Electrophysiology recording analysis

Kilosort (version 2.5)^46^ with manual curation using Phy (version 2; https://github.com/cortex-lab/phy) were used for spike sorting. A unit was considered a single unit based on manual inspection of its inter-spike interval (ISI) histogram, separation from other units and its stationarity across the session. Units that passed manual curation but had a higher ISI violation rate were called multi-units. Recording sessions were included for analysis only if they had at least 15 single units.

For the Simple Reward task, we recorded sessions from mice and targeted the ALM and tjM1. In total, 856 units (595 single units in the tjM1, and 328 single units in the ALM) were recorded in these sessions. For the Delayed Reward task, we recorded sessions from mice and targeted the ALM, tjM1, and tjS1. In total, 2011 units (934 single units in the tjM1, 704 single units in the ALM, 373 single units in the tjS1) were recorded in these sessions. For the Double Reward task, we recorded sessions from mice and targeted the ALM and tjM1. In total, 2026 units (1213 single units from the tjM1, and 813 single units from the ALM) were recorded in these sessions. For the VTA Stimulation experiments, we recorded sessions from mice and targeted the tjM1. In total, 719 single units were recorded in these sessions. For the Simple Reward Learning experiments, we recorded sessions from mice and targeted the tjM1. In total, 820 single units were recorded in these sessions. For all analyses, only single units with firing rates exceeding 0.5 Hz were included. For all analyses, spike times were binned into 5 ms bins and binned spike rates were smoothed using a causal (single-sided) gaussian kernel with standard deviation of 0.075 s.

### Linear regression and decoding

To decode behavioral kinematics from population activity, we used a time-lagged linear decoder. The decoded behavioral feature at time *t* was modeled as a weighted sum of neural activity across neurons and temporal lags:

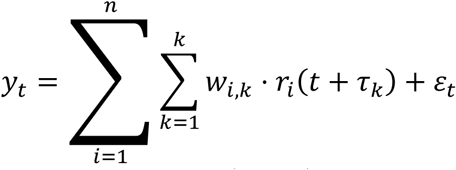

where *yₜ* is the decoded behavioral feature at time *t*, *r_i_* (*t* + 𝜏_𝑘_) is the normalized firing rate of neuron *i* at lag 𝜏_𝑘_ relative to t, 𝑤_𝑖,𝑘_ is the decoding weight for neuron *i* at lag 𝜏_𝑘_*, n* is the number of simultaneously recorded units, *k* is the number of temporal lags, and *εₜ* is the residual error at time *t*. Neural activity was included from 60 ms before to 40ms after the time point at which the behavioral feature was decoded.

To prevent overfitting, weights were estimated using ridge regression (using the MATLAB function ‘*ridge*’), which minimizes the penalized squared error:

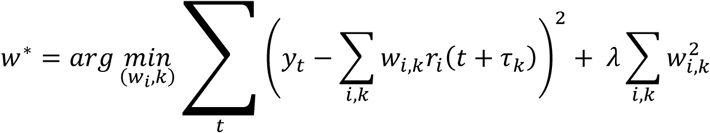

Where λ is a regularization parameter controlling the strength of the penalty on large weights.

We optimized λ by fitting the ridge regression model using only the training set trials, computed the root-mean-squared error (RMSE) between the actual and predicted tongue length and jaw position on test trials, and then selected the λ that yielded the lowest test set RMSE during the time epoch used to construct the model. We subsequently retrained the model with this optimal λ on the training set and evaluated its performance on the held-out test trials. Two separate regression models were fit: an “early” model, using the 0.3 s of data preceding the first port contact through 0.08 s following it, and a “late” model, using data from 0.25 s after the first port contact until the last contact (up to 3 s post contact). All models were evaluated on every time point within each trial. For the Simple Reward task, coefficients were learned from ∼70% of first-contact trials and tested on the remaining 30%. For the Delayed and Double Reward tasks, models were trained on ∼70% of C1-reward trials and tested on the remaining 30% of C1-reward trials as well as all C4-reward (or C1,6-reward) trials. In the VTA Stimulation task, coefficients were learned from ∼70% of trials and tested on the remaining 30%.

The normalized population spike rates were computed by z-scoring activity relative to the mean and variance of spike rates computed in one second time window before the Go Cue (from 1 second x Ntrials of single-trial data). The kinematic variables decoded included tongue length and jaw position. Features were sampled at 200 Hz. Tongue length was only calculated when the tongue was visible outside of the mouth. Consequently, time points where the tongue was not visible were set to zero. Behavioral features were min-max normalized from zero to one across all timepoints and trials.

### Decoding Index

We extracted tongue length and jaw position values across trials and corresponding model predictions from each cortical area, as well as a constant zero baseline. All data were aligned to the first lick contact and restricted to a common window (-0.08 s to 3 s). For each lick, we computed the root-mean-square error (RMSE) between all observed and predicted values across trials, pooled these errors over all sessions, and calculated the mean and 95% confidence interval of the mean. The decoding index was defined as 1 – (mean RMSE_decoded_ / mean RMSE_baseline_).

Individual licks were extracted by identifying consecutive time points during which the tongue was visible. Jaw position, unlike tongue length, is a continuous kinematic variable. To segment jaw movements into lick-aligned epochs, we identified individual jaw cycles using a trough-to-trough approach. Local minima (troughs) in the jaw position trace (as viewed from the side camera) corresponding to the minimum of jaw displacement (jaw closed), were detected, and each jaw cycle was defined as the segment of the signal between two successive troughs. This procedure captured one complete jaw movement cycle corresponding to a single lick cycle.

### Standardization of Inter Lick Intervals

Individual tongue protrusions had variable durations. Therefore, for visualization only, we time-warped data in a piecewise linear manner within each lick bout. Each tongue protrusion was stretched or compressed in time to a common value, and each inter-lick interval (ILI) was also stretched or compressed to a different, common value (such that the protrusion:ILI duration was 2:1). A lick bout was defined as a series of consecutive licks in which every ILI was shorter than 0.4 s. All quantitative analyses were obtained without time warping.

### Switching Regression Models

We used a switching regression model, also known as a generalized linear model-hidden Markov model, to identify the timepoint of disengagement in individual behavioral trials in the double reward task.

Switching regression models consist of two components: a state model and an emission model. The state model is a first-order Markov process that governs the dynamics of latent variables *z*. The emission model is a Gaussian generalized linear model with parameters β_z_ and bias *C_z_* conditioned on the latent state. We used Gaussian GLMs to relate orofacial key points *U* captured by DeepLabCut to neural principal components *X at time t*, as shown below.

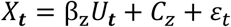

The raw data matrices were:

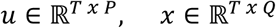

Where 𝑃=26 was the number of key point features (13 points, x and y coordinates), 𝑄=10 is the number of neural PCs, and T is the number of timepoints in a trial. The start timepoint was set 14 samples (140ms) before the GC.

For each trial, the endpoint was calculated as the last relevant lick port contact (LRC), defined as the last contact before a lapse in licking of over two times the median interlick interval for that trial. If the LRC was less than two licks after the rewarded lick, that trial was omitted from the training data.

We created design matrix ***U*** and output matrix ***X*** that included τ = 4 past time points for each sample. These matrices contain time lagged features starting 10ms before the GC up until the LRC.

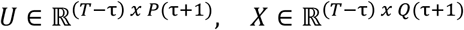

The model has parameters θ = {𝑨, π, 𝛃, 𝚺} and hyperparameter *K*, where *A* is the transition matrix, π is the initial state distribution, β and Σ are the Gaussian GLM parameters, and *K* defines the number of discrete, latent states. We selected *K* = 2, reflecting engaged and disengaged states. The initial state and transition matrices were initialized under the belief that an animal starts trials in the disengaged state and has sparse within-trial state transitions.

To initialize the engaged state’s emission model, we chose a time window where we are confident the motor cortex is engaged: movement initiation. On C1 trials, we initialized from 30ms before the first lick port contact up until the moment of contact. On C4 trials, we initialized from 30ms before the first lick port contact up to 70ms after the contact. Since we don’t have complete knowledge of when the motor cortex is disengaged during movement, we randomly initialized the disengaged state’s emission model.

To fit the switching regression model, we used Expectation-Maximization (EM). EM is a common method for finding maximum likelihood solutions to probabilistic models with latent variables. A model was fit to behavioral sessions for each brain region (tjM1 or ALM) separately. For each behavioral session - region pair, every trial was used for model training, except those filtered out by the LRC criteria. The model was fit until the log-likelihood converged to a tolerance of 10^−6^ or until 100 iterations of EM, whichever occurred first. GLM–HMM fits produced quantitatively similar results for tjM1 and ALM. As a result, data from these two motor cortical areas were combined, and Fig. 2j–m shows pooled results from tjM1 and ALM.

We utilized StateSpaceDynamics.jl, a Julia language package for state space modeling. After model fitting, we performed state inference using the Forward-Backward algorithm implementation in StateSpaceDynamics.jl.

### Statistics and reproducibility

No statistical methods were used to predetermine sample sizes. All *t*-tests were two-sided unless stated otherwise. In Fig. 3h and EDFig. 4j, to evaluate whether the time evolution of decoding accuracy was different across tasks/days, we tested the change in decoding accuracy for each lick relative to decoding accuracy for the first lick, rather than the absolute decoding accuracy, as baseline decoding accuracy varied somewhat across sessions/individuals. All animals from which electrophysiology and/or behavioral data were collected were included in the study. Trial type, and the trials on which optogenetic perturbations were applied, were randomly determined during electrophysiology and behavioral data collection. Experimenters were blinded to trial type during all data collection and spike sorting.

---

**Extended Data Figure 1 -.**
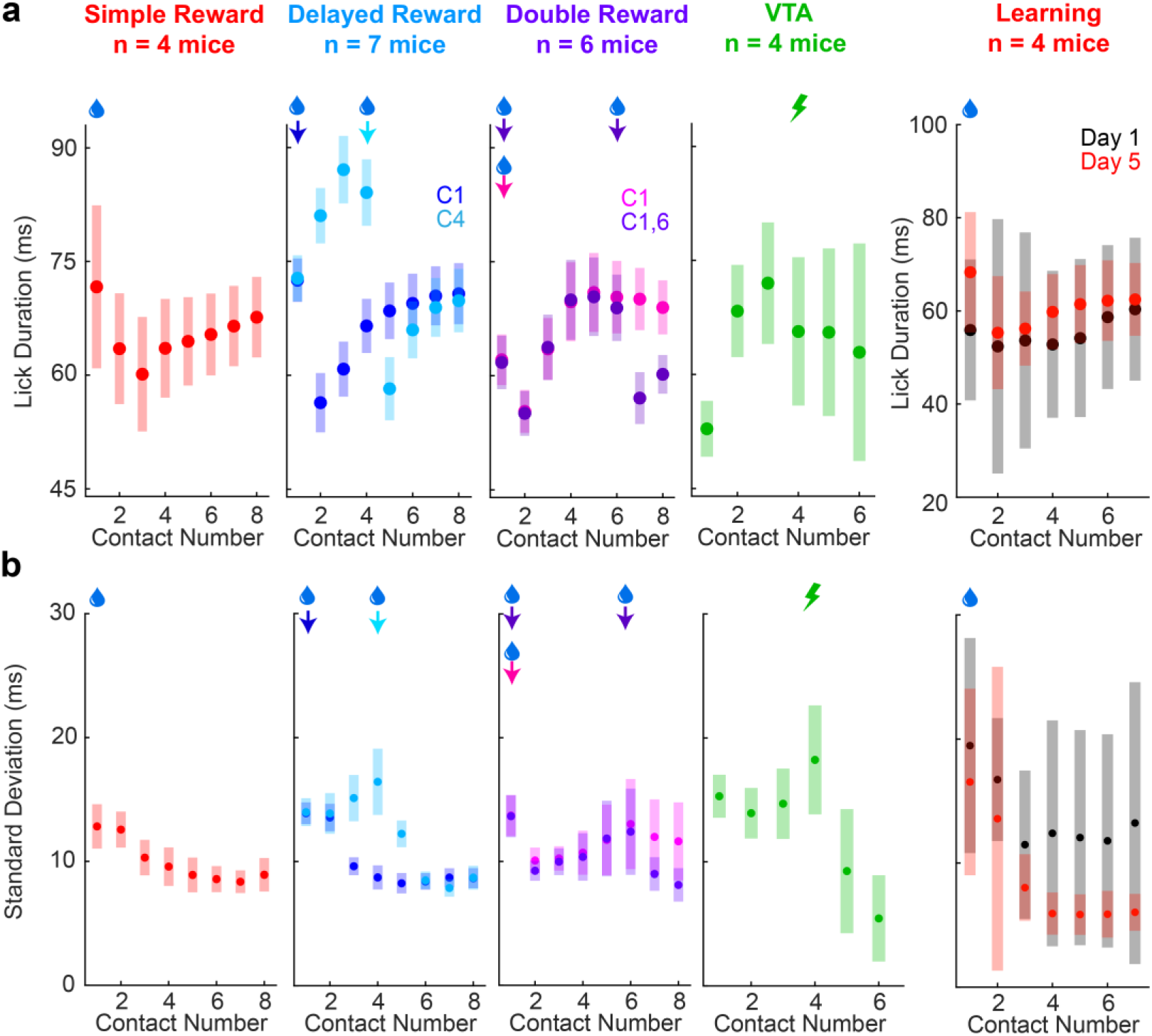
Tongue kinematics across tasks. **a.** Mean duration of tongue protrusions across tasks. **b.** Standard deviation of tongue protrusion duration across tasks. Shaded regions and error bars indicate 95% confidence intervals in all panels.

**Extended Data Figure 2 -.**
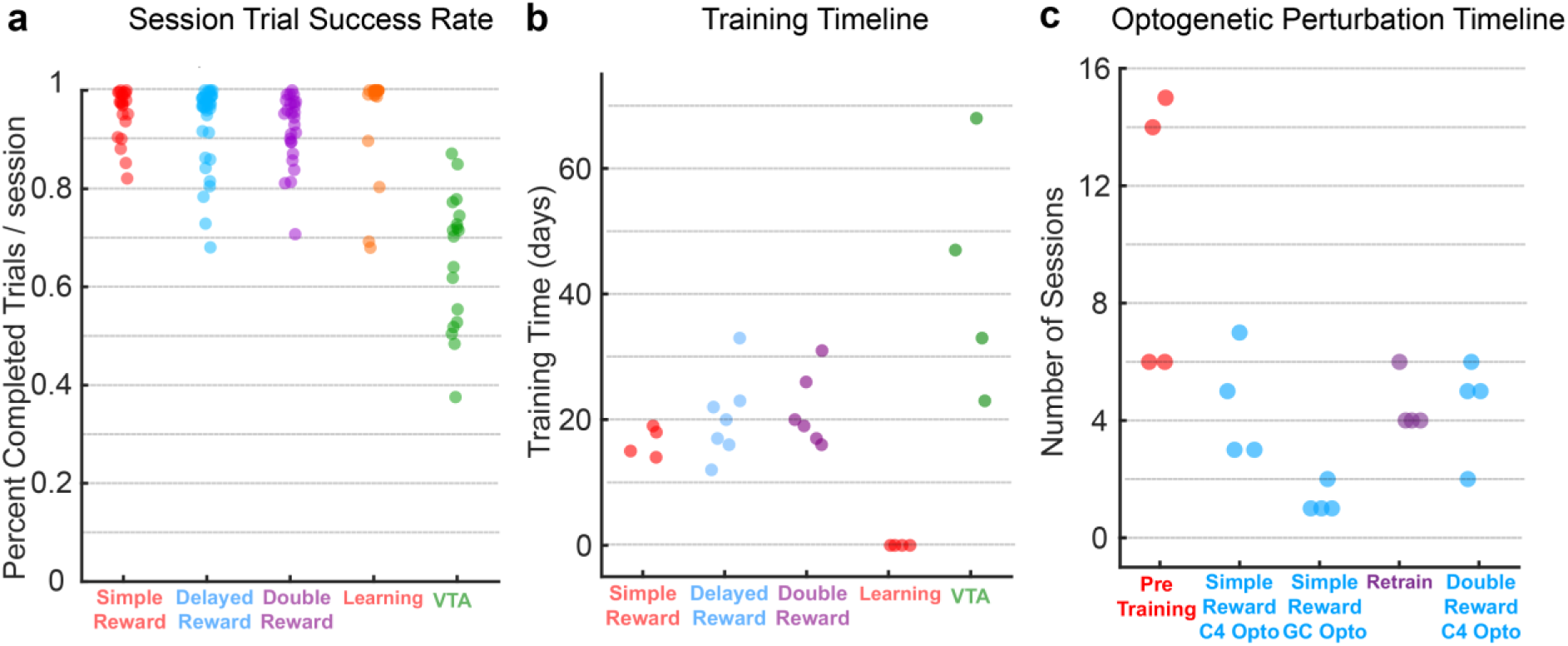
Session statistics and experiment timelines. **a.** Proportion of trials successfully completed in each session across tasks. **b.** Number of training sessions prior to the first recording session across tasks. **c.** Timeline of optogenetic perturbation experiments indicating the number of sessions each animal performed in each experimental epoch.

**Extended Data Figure 3 -.**
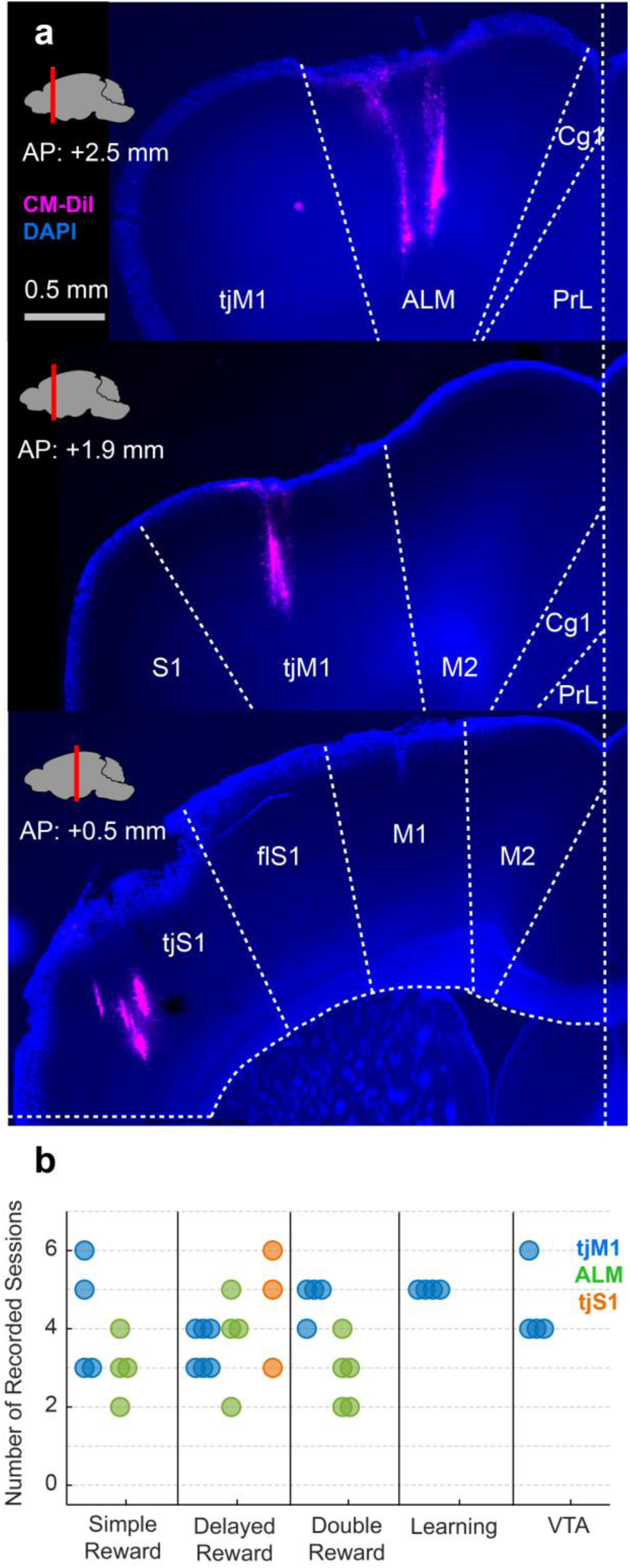
Probe tract histology and summary of experiments. **a.** Example images showing DAPI (blue) and CM-Dil (magenta) fluorescence in the ALM (*top*), tjM1 (*middle*), and tjS1 (*bottom*). **b.** Number of sessions in which electrophysiological recordings were conducted for each animal across regions and tasks.

**Extended Data Figure 4 -.**
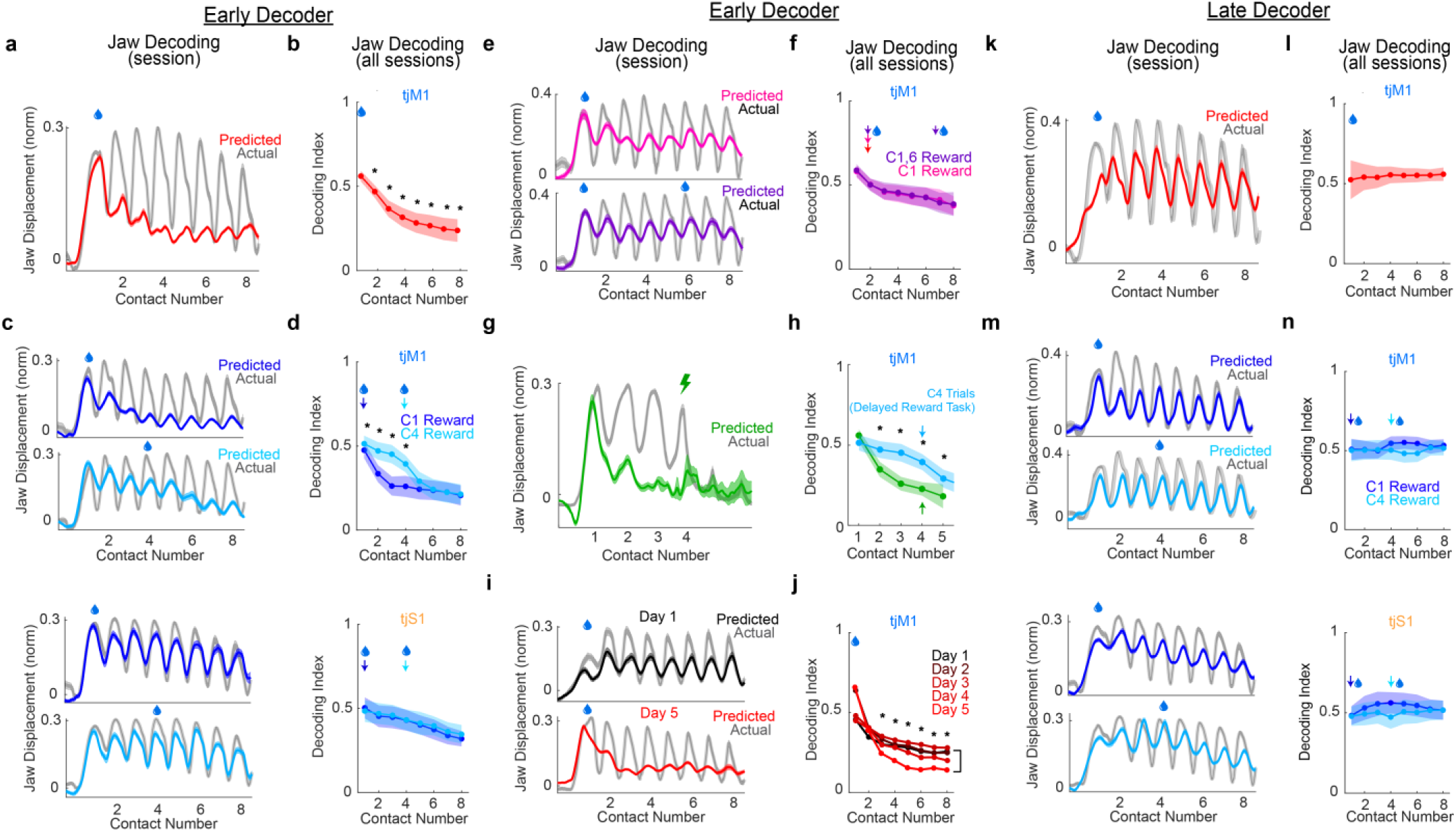
Jaw position decoding. **a.** Jaw position decoder trained on ‘early’ activity (during the first protrusion) in the tjM1 in an example Simple Reward Task session. **b.** Decoding index (normalized model RMSE, see ***Methods***) across all sessions (asterisks: p<0.005 comparing subsequent protrusions to the first, one-tailed t-test). **c, d.** Same as (a) and (b) but for the tjM1 (*top*) and tjS1 (*bottom*) in the Delayed Reward Task (asterisks: p<0.005 comparing trial types, two-tailed t-test). **e,f.** Same as (a) and (b) but for the Double Reward Task. **g,h.** Same as (a) and (b) but for the VTA Stimulation Task (asterisks: p<0.005 comparing C4-reward trials in the VTA Stimulation Task (*green*) and Delayed Reward Task (*blue*); one-tailed t-test). **i,j.** Same as (a) and (b) but for each consecutive day of training during task acquisition (asterisks: p<0.005 comparing Day 1 and Day 5, one-tailed t-test). **k,l.** Same as (a) and (b) but for a Jaw position decoder trained on ‘late’ activity (subsequent protrusions after the first; see ***Methods*). m,n.** Same as (c) and (d) but for the late decoder. Shaded regions and error bars indicate 95% confidence intervals in all panels.

**Extended Data Figure 5 -.**
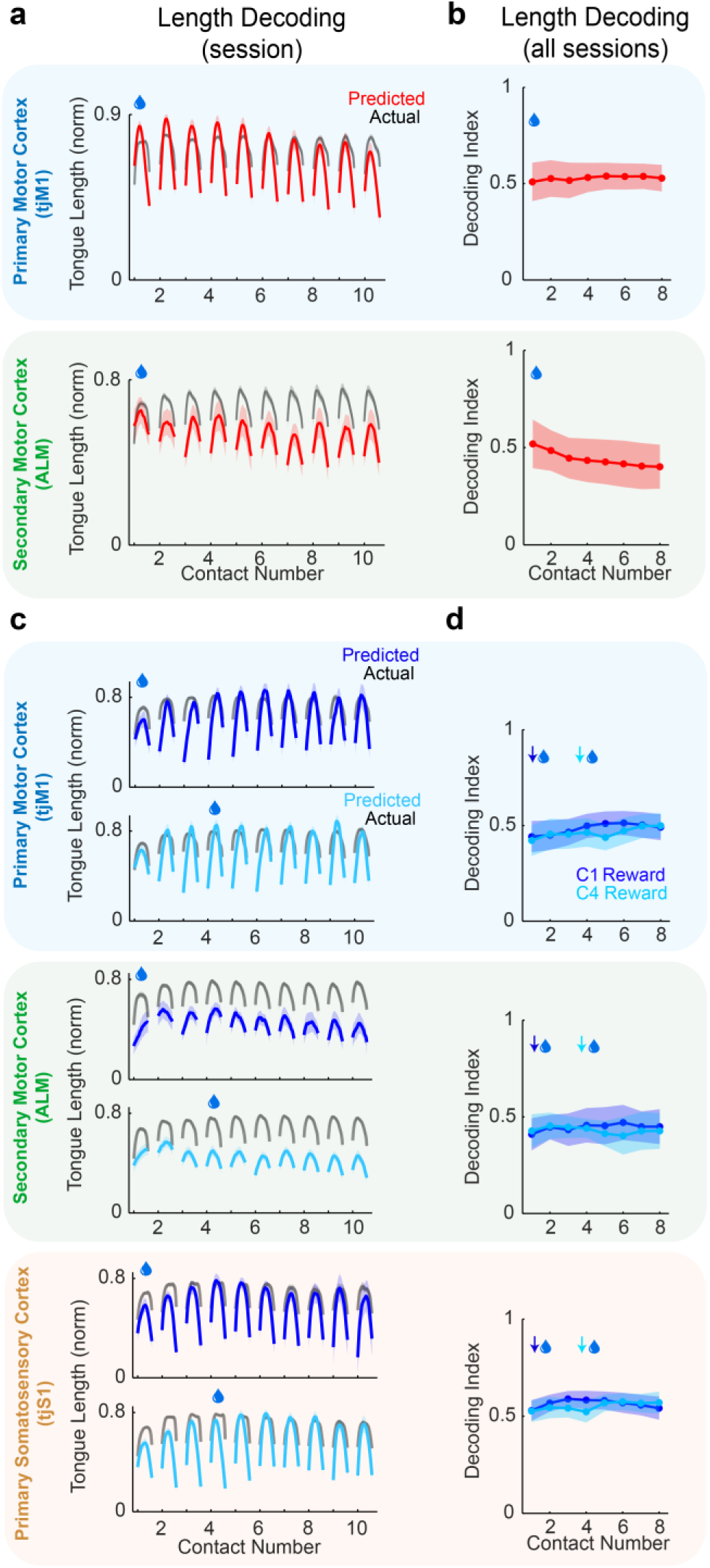
Tongue length predictions from ‘Late’ activity. **a.** Tongue length decoder trained on ‘late’ activity (subsequent protrusions after the first on C_1_-reward trials; see ***Methods***) from the tjM1(*top*) and ALM (*bottom*) in an example Simple Reward Task session. **b.** ‘Decoding index’ (normalized model RMSE, see ***Methods***) across all sessions (asterisks: p<0.005 comparing subsequent protrusions to the first, one-tailed t-test). **c,d.** Same as (a) and (b) but for the tjM1 (*top*), ALM (*middle*), and tjS1 (*bottom*) in the Delayed Reward Task. Shaded regions and error bars indicate 95% confidence intervals in all panels.

**Extended Data Figure 6 -.**
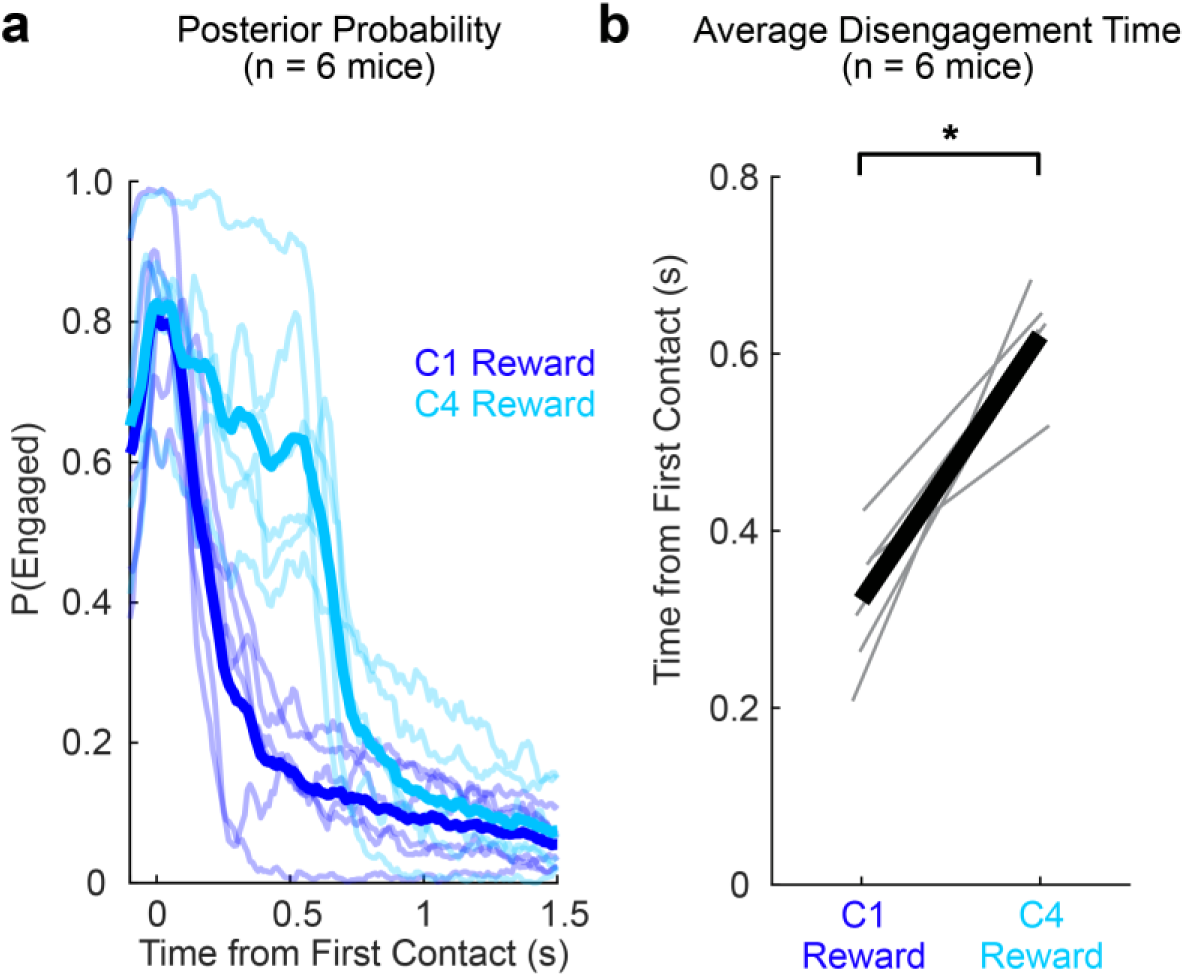
HMM-GLM results across individuals. **a.** Posterior probability of the engaged state for C1- (*dark blue*) and C4-reward trials (*light blue*) trials. Inferences aligned to the first port contact (C1). Individual animals (*thin lines*) and across-animal averages (*thick lines*) shown. **b.** Average disengagement time relative to the first port contact on C1- and C4-reward trials for individual animals (*light lines*) and across-animal averages (*dark lines*). Disengagement times were significantly later C4-reward trials compared to C1-reward trials (asterisk; p<0.005, one-tailed t-test).

**Extended Data Figure 7 -.**
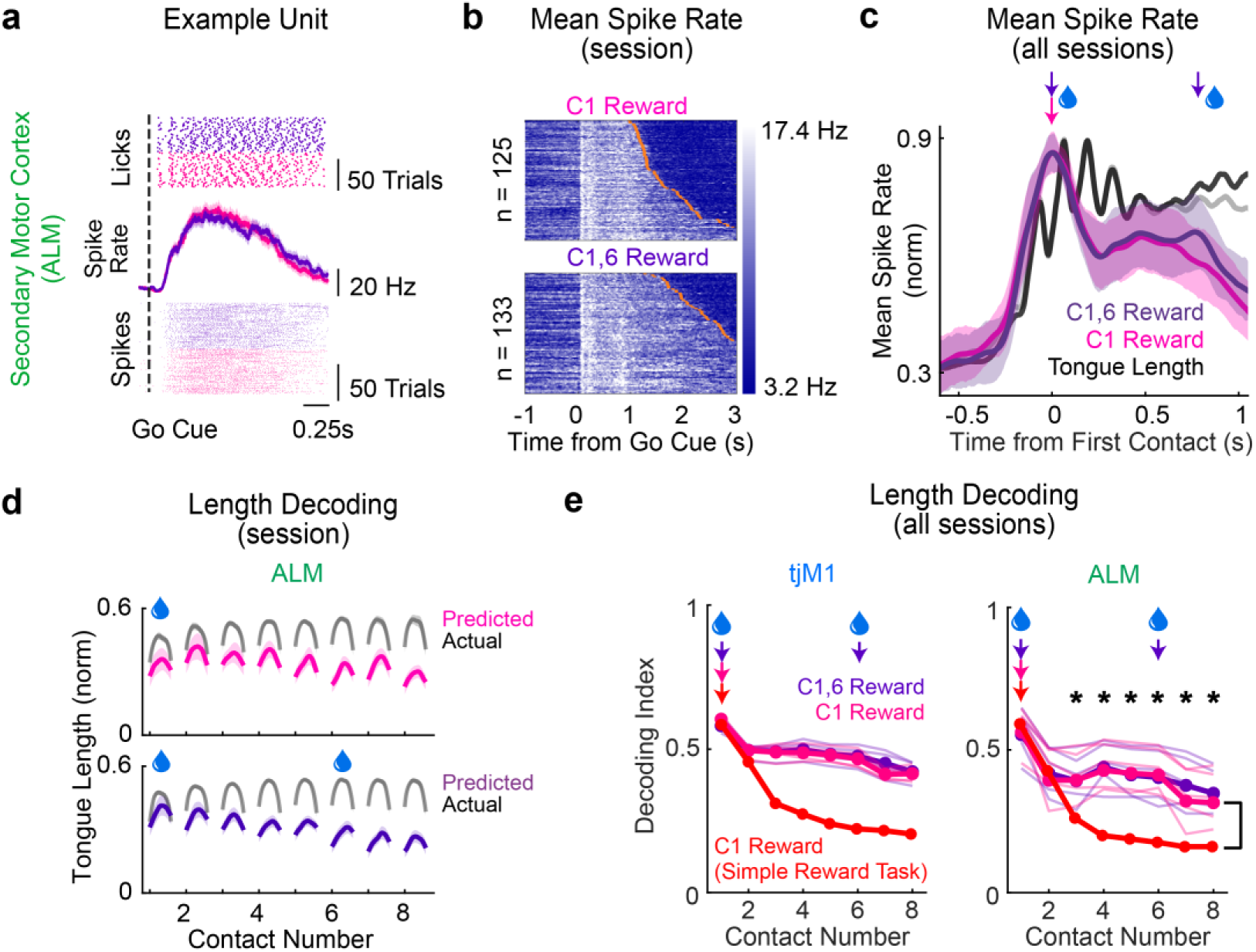
ALM dynamics in the Double Reward Task. **a.** Rasters of port contacts (*top*) and spikes (*middle*) and trial-averaged spike rates (*bottom)* for example units in the ALM. **b.** Mean spike rate across all units in the ALM in an example session. Trials sorted by the timing of the last lick in a bout (*orange ticks*). **c.** Mean session-normalized spike rates on each trial type in the ALM across all sessions (n = 14 sessions, 4 animals). Mean tongue length on C1-reward trials (*gray*) and C1,6-reward trials (*black*) indicated for reference. **d.** Tongue length predictions from the ALM in an example session. **e.** Decoding index (see ***Methods***) across all sessions (asterisks: p<0.005 comparing C1-reward trials in the Simple Reward Task and Double Reward Task; two-tailed t-test) for the tjM1 and ALM (n = 14 sessions, 4 animals). Results for individual animals (*thin lines*; Double Reward Task only for clarity) and across-animal averages (*thick lines*) shown. Shaded regions and error bars indicate 95% confidence intervals in all panels.

**Extended Data Figure 8 -.**
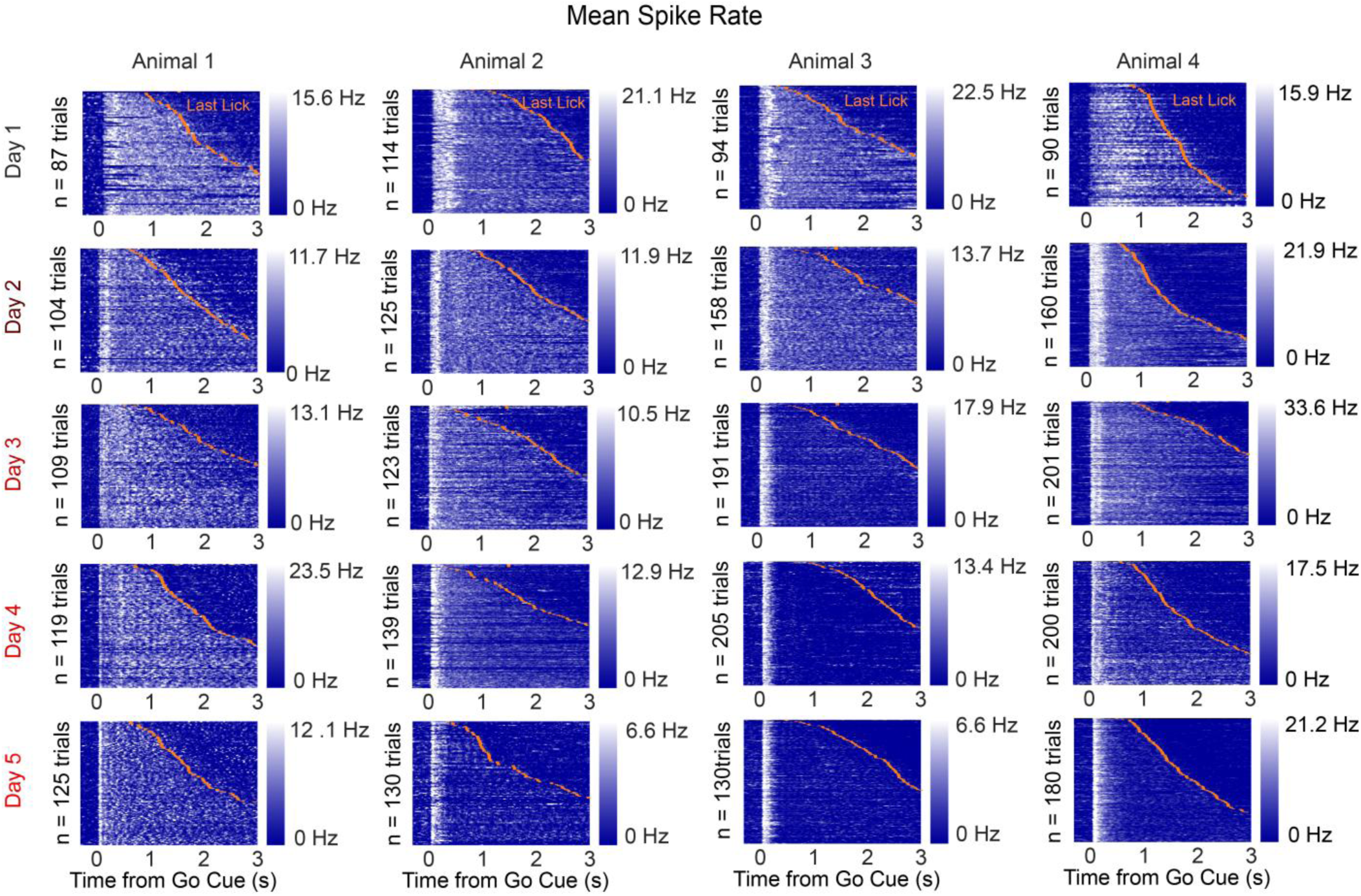
Neural activity during task acquisition. Baseline-subtracted mean spike rate, averaged across all units in the tjM1, during the first five consecutive days of training. Trials are sorted by the timing of the last lick in a bout (*orange ticks*).

## SUPPLEMENTARY VIDEOS

**Supplementary Video 1 – Motor cortex photoinhibition at the fourth port contact in the Simple Reward Task.** Three example optogenetic photoinhibition (*left*) and control (*right*) trials, all from the same animal, are displayed with the photoinhibition period indicated by a white square in the bottom-left corner.

**Supplementary Video 2 – Motor cortex photoinhibition at the Go Cue in the Simple Reward Task.** Same as Supplementary Video 1, but with photoinhibition beginning at the Go Cue.

**Supplementary Video 3 – Example C4 trials in the Delayed Reward Task.** Two example C4 trials are shown with the tip of the tongue labeled with blue dots during the period before reward and red dots following reward delivery.

**Supplementary Video 4 – Motor cortex photoinhibition at the fourth port contact in the Double Reward Task.** Same as Supplementary Video 1 except animal was performing the Double Reward Task instead of the Simple Reward Task. Same animal as Supplementary Video 1.

## Notes

### Competing Interest Statement

The authors have declared no competing interest.

